# Inverse identification of region-specific hyperelastic material parameters for human brain tissue

**DOI:** 10.1101/2022.12.19.521022

**Authors:** Jan Hinrichsen, Nina Reiter, Lars Bräuer, Friedrich Paulsen, Stefan Kaessmair, Silvia Budday

## Abstract

The identification of material parameters accurately describing the region-dependent mechanical behavior of human brain tissue is crucial for computational models used to assist, e.g., the development of safety equipment like helmets or the planning and execution of brain surgery. While the division of the human brain into different anatomical regions is well established, knowledge about regions with distinct mechanical properties remains limited. Here, we establish an inverse parameter identification scheme using a hyperelastic Ogden model and experimental data from multi-modal testing of tissue from 19 anatomical human brain regions to identify mechanically distinct regions and provide the corresponding material parameters. We assign the 19 anatomical regions to nine governing regions based on similar parameters and microstructures. Statistical analyses confirm differences between the regions and indicate that at least the corpus callosum and the corona radiata should be assigned different material parameters in computational models of the human brain. We provide a total of four parameter sets based on the two initial Poisson’s ratios of 0.45 and 0.49 as well as the pre- and unconditioned experimental responses, respectively. Our results highlight the close interrelation between the Poisson’s ratio and the remaining model parameters.

The identified parameters will contribute to more precise computational models enabling spatially resolved predictions of the stress and strain states in human brains under complex mechanical loading conditions.

## Introduction

As powerful computational resources have become available, it is now feasible to run complex mechanical simulation models of the brain. An emerging application of mechanical models for brain tissue is the simulation of neurosurgeries that enable surgeons to learn the procedures ’in the dry’ without any risk of harming the patient (Sase et al., 2015; Hansen et al., 2004; Delorme et al., 2012). Another application is the design of protective equipment. Brain injuries are a major health issue. A study conducted by Majdan et al., 2017 reported a total number of 17.049 deaths related to traumatic brain injury (TBI) in 16 european countries during the year 2013. The development of new as well as the improvement of existing protective measures like helmets can contribute to reduce this number. An overview of currently used brain biomechanical models is given in the review by Ji et al., 2022. Besides the simulation of external loads in the aforementioned scenarios, there are also models predicting phenomena linked to internal processes, such as (abnormal) cortical folding during brain development (Garcia et al., 2018), associated cellular processes (Budday et al., 2015; Zarzor et al., 2021) and diseases like epilepsy (Blumcke et al., 2021) or Alzheimer’s (Weickenmeier et al., 2018).

A key prerequisite for accurate simulation models are constitutive relations to solve the underlying boundary value problem arising from Cauchy’s equation of motion. Constitutive equations themselves require the identification of appropriate material parameters. Their ability to approximate the actual mechanical behavior has a significant influence on the accuracy of the output of the whole model. Most of the currently used models assume homogeneous material properties for brain tissue as a whole (Zong et al., 2006; Yan et al., 2011; Fernandes et al., 2018; Horgan et al., 2003; Kang et al., 1997). Only some distinguish between brain and brainstem (Ho et al., 2009; Ghajari et al., 2017) or between gray and white matter (Mao et al., 2013). Other approaches have incorporated a more heterogeneous distribution of material properties, for example by utilizing data from magnet resonance elastography (MRE) (Zhao et al., 2022; Giudice et al., 2021), or have accounted for axon fiber tracts (Wu et al., 2019; Li et al., 2021). It is commonly assumed that brain tissue is nearly incompressible, which is often implemented in terms of a compressible formulation with a high bulk modulus in finite element models (Zong et al., 2006; Yan et al., 2011; Fernandes et al., 2018; Kang et al., 1997). Different values for the initial value of the Poisson’s ratio *v* have been used for such simulations, e.g., 0.49995 (MacManus et al., 2018), 0.495 (Pierrat et al., 2018), and 0.49 (Shafieian et al., 2009).

There have been numerous experimental studies characterizing the mechanical behavior of brain tissue. An overview of experiments is given in the review by Faber et al., 2022. Material parameters can be identified by fitting a mechanical model with an underlying parameterized constitutive relation (a material model) to the experimental data. Budday et al., 2017a showed the importance of simultaneously considering multiple loading modes when aiming to calibrate reliable material parameters. Due to the observed heterogeneity in mechanical properties, it is also desirable to test each specimen under all loading modes as opposed to using separate samples for the different tested modes. Although there have been more experiments on human brain tissue in recent years (Karimi et al., 2019; Forte et al., 2017; Jin et al., 2013; Chatelin et al., 2012; Zhu et al., 2010; Jin et al., 2013; MacManus et al., 2020; Finan et al., 2017; Greiner et al., 2021; Budday et al., 2017a), there are still only a few testing human brains under multiple loading modes (Budday et al., 2017a; Greiner et al., 2021; Jin et al., 2013). There are also significant differences regarding the spatial resolution of experimental studies, i.e., the classification into different regions. While some report experimental results for gray and white matter (Forte et al., 2017), others differentiate into three regions, i.e., corona radiata, thalamus and brainstem (Chatelin et al., 2012), or up to four regions, i.e., corpus callosum, corona radiata, basal ganglia and cortex (Budday et al., 2017a). The distinction of these regions has so far been based on anatomical knowledge. However, to what extent anatomically distinct regions coincide with mechanically distinct regions remains largely unknown.

Most material parameters for brain tissue have been identified based on the assumption of homogeneous deformation states during experiments (Budday et al., 2017a; Laksari et al., 2012; Mihai et al., 2017a). In experimental tests other than compression, however, the specimens need to be fixed to the specimen holders, which is usually achieved by gluing. The thereby introduced non-slipping boundary conditions lead to an inhomogeneous deformation state. By running finite element simulations with realistic boundary conditions using parameters obtained based on the assumption of a homogeneous deformation, it has been shown that this assumption introduces a notable error in the model predictions (Budday et al., 2019; Felfelian et al., 2019; Voyiadjis et al., 2018).

In this paper, we aim to identify mechanically distinct regions that should be accounted for in finite element models of the human brain and provide corresponding hyperelastic material parameters. To this end, we test human brain tissue extracted from a total of 19 anatomical regions under cyclic compression, tension, as well as torsional shear. To capture the inhomogeneous deformation state during testing caused by non-slip boundary conditions, we implement a quasistatic finite element model that simulates the testing procedure. We use a hyperelastic one-term Ogden model based on previous results of different groups who successfully used Ogden-type models to model the mechanical behavior of brain tissue (Mihai et al., 2015; Miller et al., 2000; Budday et al., 2017a; Hosseini-Farid et al., 2019). By coupling our simulation model with an optimization routine, we identify the optimal set of material parameters. We fit parameters for the first and third loading cycle separately to represent the un- and preconditioned response, respectively. Furthermore, we used two different initial Poisson’s ratios of 0.45 and 0.49 to compare how a different assumed compressibility affects the results.

## Methods

### Human brains

We obtained seven whole human brains including the cerebrum, cerebellum and brainstem from three female and four male body donors who had given their written consent to donate their body to research. Table 1 gives an overview of the obtained brains including the age of the respective body donor and the cause of death. None of the body donors had suffered from any neurological disease known to affect the microstructure of the brain. We note that we could not find metastases in the brain from donors 3, 5, and 6, who had died from cancer. In brain number 7, however, we found one metastasis in the left cerebellar peduncle. The remaining tissue did not show any visible abnormalities. The brains 1-3 and 5-6 were immersed in cerebrospinal fluid surrogate (CSFS) during transport. Brain 4 was kept in phosphate buffered saline solution (PBS) and brain 7 in Ringer’s solution, as the otherwise preferred CSFS was not available on short notice. CSFS closely matches the electrolyte concentrations of cerebrospinal fluid and is prepared from high purity water and analytical grade reagents. The constituents of all three solutions are listed in Table 2. We received the brains between 9 and 24 h *post mortem* and directly cut them into 1 cm thick coronal slices that we kept refrigerated at 4°C in the solution they had arrived in (CSFS, PBS, or Ringer’s solution) until mechanical testing. The mechanical experiments were completed within 72 hours *post mortem.*

**Table 1.** Human brains.

| Brain | Sex | Age | Cause of death |
| --- | --- | --- | --- |
| 1 | male | 92 | cardiovascular failure |
| 2 | female | 62 | liver and kidney failure |
| 3 | male | 68 | metastasizing bronchial carcinoma |
| 4 | male | 75 | cardiac insufficiency |
| 5 | male | 75 | metastasizing bronchial carcinoma |
| 6 | female | 77 | metastasizing pancreas carcinoma |
| 7 | female | 69 | metastasizing breast cancer |

**Table 2.** Storage solutions.

| Solutions | Constituents (in mM/100ml) |
| --- | --- |
| CSFS | Na+ 150, K+ 3.0, Ca2+1.4, Mg2+ 0.8, P 1.0, Cl- 155 |
| PBS | NaCl 137, KCl 2.7, Phosphate buffer 11.9 |
| Ringer's | NaCl 1.24×1e-4, NaH2PO4 1.25, MgSO4 1.8, CaCl2 1.6, KCl 3.0, Glucose 10 |

### Specimen preparation

We used a biopsy punch to extract cylindrical samples of 8mm diameter, as shown in Figure 2a and b. To ensure that the specimens only experienced small deformations before being probed mechanically, we punched them out of coronal slices while the slices were immersed in CSFS, PBS, or Ringer’s solution. Like this, the cylindrical specimens could slide out of the biopsy punch without adhering to it. If the small cylinders had a height of more than 6 mm, we carefully shortened them with a surgical scalpel. The final specimen height as recorded by the rheometer (see also Figure) varied between 2.7mm and 7.2mm with a mean of 4.9mm. For most regions, it was possible to extract homogeneous specimens of this size. Exceptions are brain regions that contain both white and gray matter tissue, such as the midbrain, pons, and medulla. Samples of the deep cerebellar nuclei contained a certain amount of cerebellar white matter because the cerebellar nuclei are too small to be probed separately. Figure 1 gives an overview of samples extracted from each brain region. The corresponding abbreviations are introduced in Table 3.

**Table 3.**
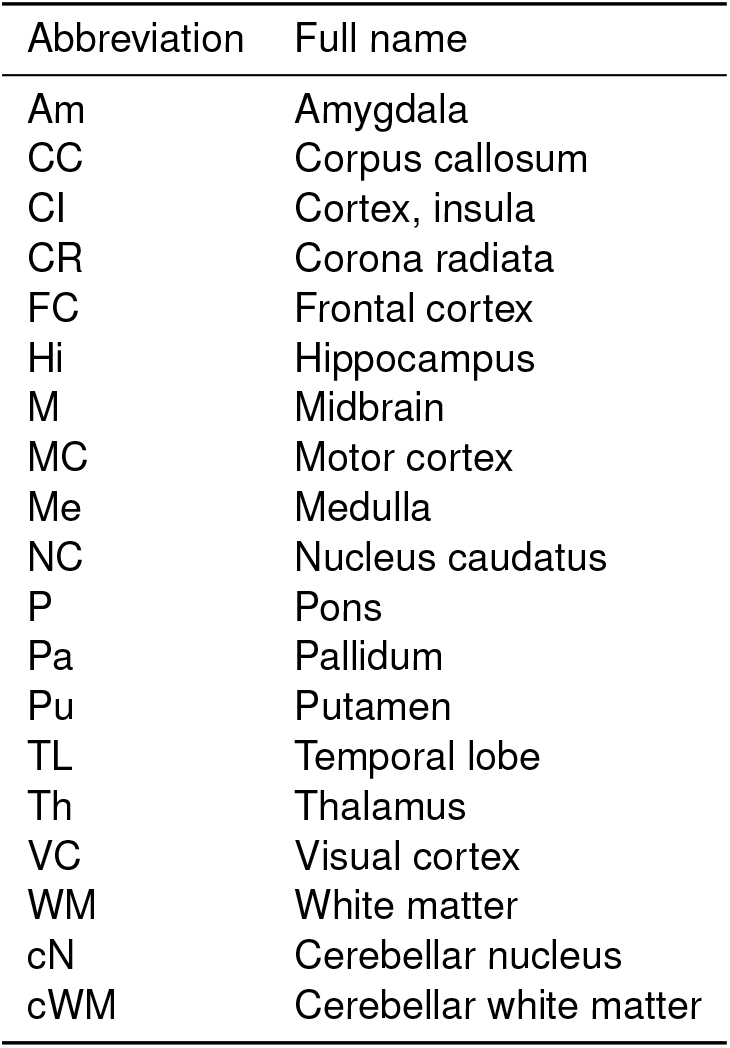
Regions and their abbreviations.

**Figure 1.**
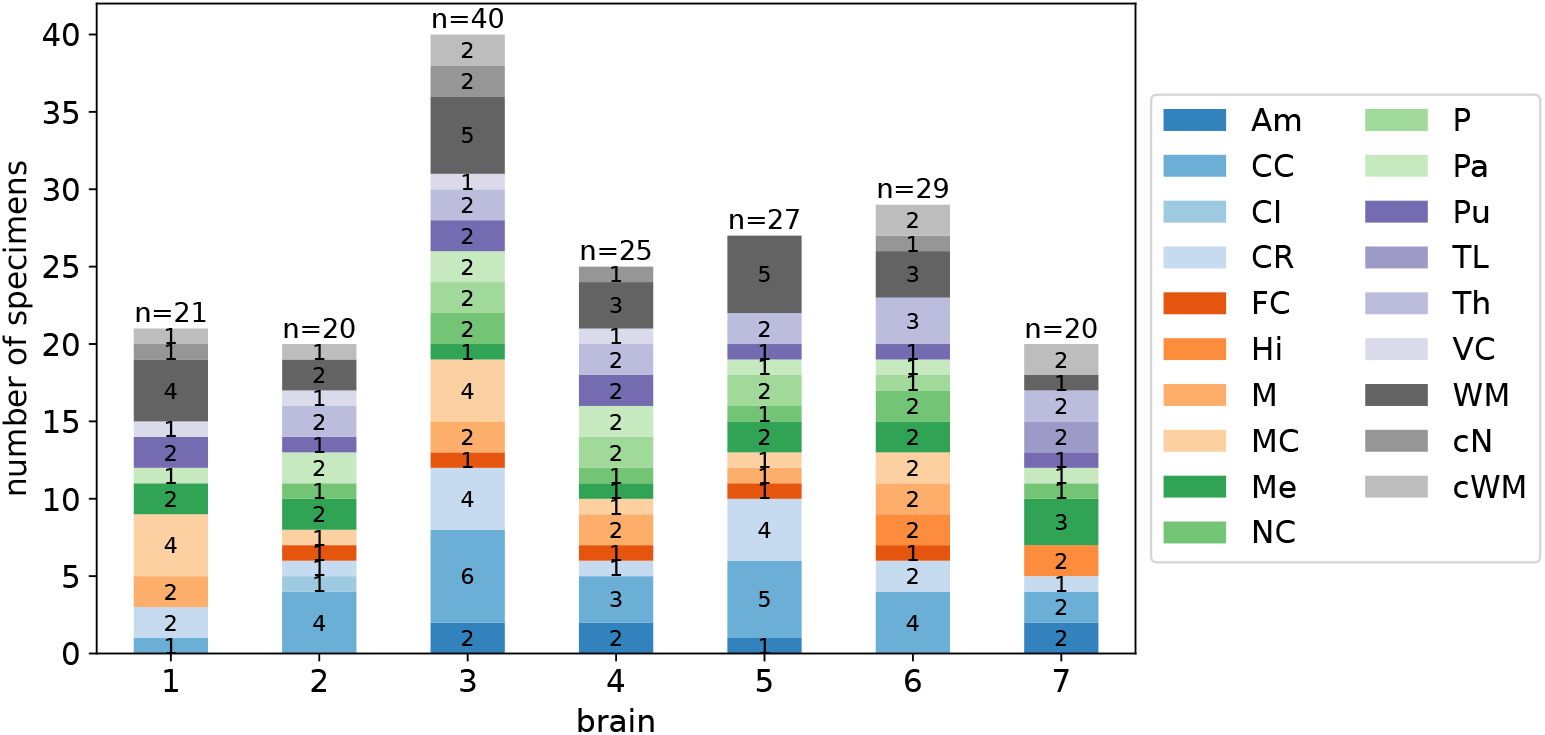
Number of specimens extracted from the different regions in the individual brains.

**Figure 2.**
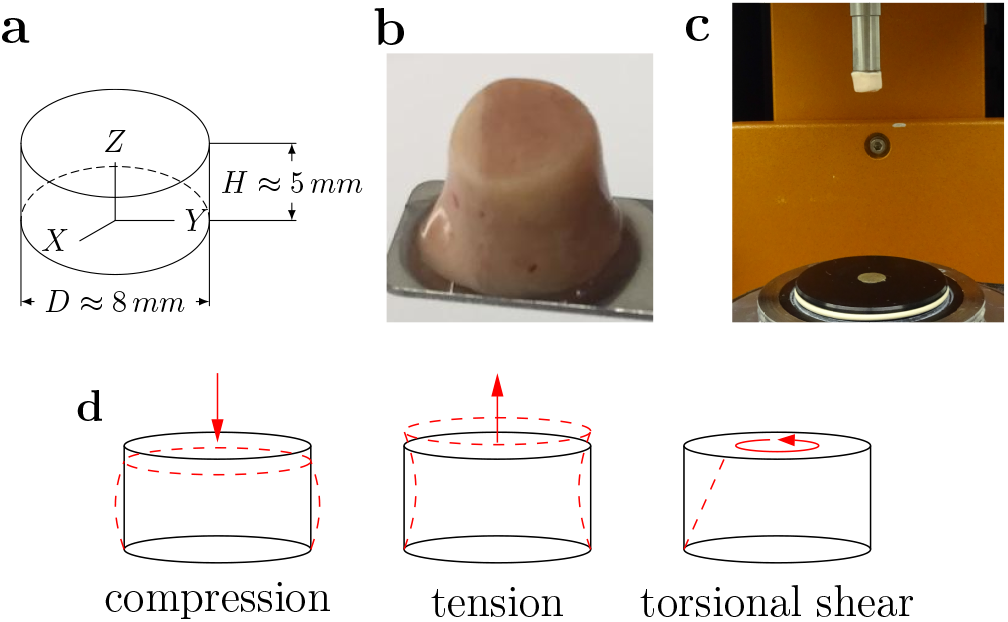
Specimen preparation and experimental setup. (**a**) Idealized geometry and dimensions of the specimens. (**b**) Extracted brain tissue specimen prepared for testing. (**c**) Specimen glued to the upper specimen holder of the rheometer. (**d**) Tested loading modes. Adapted from Faber et al., 2022.

### Experimental setup

We used a Discovery HR-3 rheometer from TA instruments (New Castle, Delaware, USA) to measure the tissue response under compression, tension, and torsional shear (see Figure 2c and d). After gluing sandpaper to the specimen holders and calibrating the instrument, we fixed the specimens to the sandpaper on the upper and lower specimen holders using superglue. We waited 30 to 60 s to let the glue dry before adding PBS to immerse the specimen and keep it hydrated during the experiment. We conducted all tests at 37°C. Figure 2c shows a specimen that has been fixed to the upper specimen holder. The testing protocol is summarized in Table 4. We first applied three cycles of compression and tension with a loading velocity of 40 μm/s and minimum and maximum stretches of *λ* = [*H* + Δ*z*]/*H* = 0.85 and *λ* = 1.15, where *H* denotes the initial specimen height and Δ*z* the displacement in the direction of loading. Subsequently, we performed a compression relaxation test at *λ* = 0.85 with a loading velocity of 100 μm/s and a holding period of 300 s, and a tension relaxation test at *λ* = 1.15, with the same loading velocity and holding period. Then, we performed two sets of cyclic torsional shear tests with three cycles and a maximum shear strain of *γ* = 0.15 and *γ* = 0.3, respectively. For compression and tension tests, we recorded the corresponding force *f_z_* and determined the nominal stress as *P_exp_* = *f_z_/A*, where *A* is the cross-sectional area of the specimen in the undeformed configuration. For torsional shear tests, we recorded the corresponding torque *t* and determined the torsional shear stress as *τ* = 2*t/πr*^3^.

**Table 4.** Testing protocol.

|  |  |  |
| --- | --- | --- |
| 1 | Frequency sweep | at 1 % strain |
| 2 | Cyclic compression/tension in z-direction | up to 15 % strain |
| 3 | Compression relaxation in z-direction | at 15 % strain; hold time=300 s |
| 4 | Tension relaxation in z-direction | at 15 % strain; hold time=300 s |
| 5 | Cyclic torsional shear | 3 Cycles up to an amount of shear of $\gamma = 0.15$ |
| 6 | Cyclic torsional shear | 3 Cycles up to an amount of shear of $\gamma = 0.3$ |

### Finite element model

#### Boundary value problem

The balance of linear momentum in a quasistatic setting can be written as div***σ*** + **b** = **0**, where ***σ*** denotes the Cauchy stress tensor and **b** the vector of volume forces. The deformation map *φ*(**X**) maps the tissue from the undeformed (material) configuration **X** to the deformed configuration **x**. To describe the deformation of our specimens during testing, we introduce the deformation gradient as **F** = ∇_**X***φ*_(**X**), which maps line elements from the material configuration *X* to the spatial configuration *x*. **F** can be uniquely decomposed into a rotation tensor **R** and a stretch tensor **v** so that **F** = **vR** with **R**^T^**R** = **I** and **v** = **v**^T^. The eigenvalues of **v** are identified as the principal stretches *λ_a_* and can be obtained as the square root of the eigenvalues of the left Cauchy-Green strain tensor **b** = **FF**^*T*^. Our goal is to find the displacement field **u**, defined as **u** = **x** – **X**, that solves div***σ*** + **b** = **0** on the region Ω, representing our specimen. The boundary surface *∂*Ω can be split into a part *∂*Ω_*u*_, where we prescribe the displacements as Dirichlet boundary conditions **u** = **ū**, and a part *∂*Ω_*σ*_, where we prescribe the tractions as Neumann boundary conditions 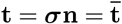. As we neglect the influence of volume forces, we set **b** = **0** and obtain the boundary value problem in its strong form

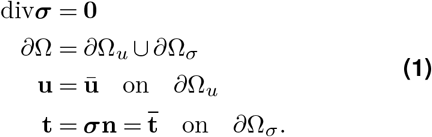

After reformulation in its weak form and a subsequent linearization, we use the finite element method (FEM) to solve this set of equations. The high strain values (up to 15% nominal strain) applied in the experiments in conjunction with the nonlinear stress-strain relation of the used constitutive model results in a highly nonlinear problem. Therefore, we employ an iterative solution scheme based on the Newton-Raphson method. We implement the finite element model using the open source finite element library deal.ii (Arndt et al., 2021).

#### Hyperelastic constitutive model: modified one-term Ogden model

To solve the boundary value problem (Equation 1), we need to specify the constitutive equation relating displacements **u** to the stresses ***σ*** (Nair, 2009). In this study, we neglect time-dependent effects and focus on the time-independent, hyperelastic material response of brain tissue. In this case, the stress response depends only on the deformation state. To characterize the constitutive behavior of a hyperelastic material, a strain energy function Ψ is defined. It is then possible to obtain from the second law of thermodynamics (Holzapfel, 2000) the relation

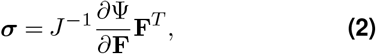

where *J* denotes the volume ratio *J* = det**F**. A common approach is to split the strain energy function into an isochoric and a volumetric part

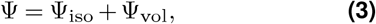

based on the assumption that the material behaves different in shear and bulk. Based on our previous works (Budday et al., 2017a; Budday et al., 2019), we use a reformulated version of the Ogden model (Ogden, 1972) in terms of the shear modulus *μ* for the isochoric part

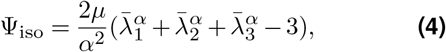

with the isochoric principal stretches 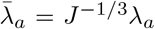, and the nonlinearity parameter *α*. The one-term Ogden model achieved promising results when fitting an incompressible analytical implementation of the model simultaneously to compression, tension, and shear experimental data (Budday et al., 2017a). In particular, it is capable of capturing the pronounced compression-tension asymmetry observed in experiments.

By applying the chain rule, it is possible to write Equation 2 in terms of the principal stretches

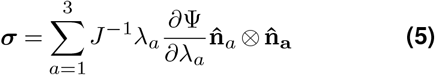

which can be interpreted as spectral form of ***σ*** with the principal values 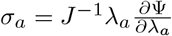 and the eigenvectors of the left Cauchy-Green tensor 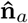. For the principal values of the isochoric stress *σ*_iso *α*_, we obtain

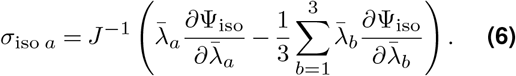

For the volumetric part of the strain energy, we choose a formulation proposed by Ogden, 1972,

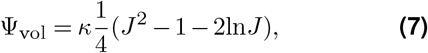

where the empirical coefficient in the original formulation has been set to 2, as introduced by Simo et al., 1992. The bulk modulus *κ* characterizes the resistance of the material against volume changes and is here calculated from the shear modulus and the initial Poisson’s ratio *v* through the relation

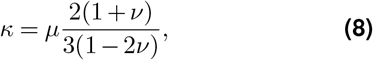

taken from the linear elastic regime. We note that we use this initial Poisson’s ratio in this context to obtain feasible estimates for the bulk modulus *κ*, where *v* serves as a measure of the compressiblity with *v* = 0.5 representing the incompressible limit resulting in lim_*v*→0.5_*κ* = ∞. While as per definition in the finite regime, a Poisson’s ratio of *v* = 0.5 is not necessarily indicating incompressible behavior (Voyiadjis et al., 2018), we here treat it as a constant rather than a function depending on the chosen strain measure and the current deformation. A constant Poisson’s ratio is often used in the literature in the context of parameter identification for brain tissue (MacManus et al., 2018; Pierrat et al., 2018; Shafieian et al., 2009; Hosseini-Farid et al., 2019). Still, it has to be noted that the interpretation of its meaning remains questionable when it is applied to different constitutive formulations. A comprehensive overview of constitutive parameters, including the Poisson’s ratio, in the context of finite elasticity and their relation to the partly available linear counterparts is given by Mihai et al., 2017b. In our case, we relate *κ*to the shear modulus by Equation 8, as the available experimental data does not contain information about the volumetric deformation of the specimens and we can, therefore, not expect to obtain reasonable values when fitting *κ* to the data.

As it is common for soft tissues, brain tissue is often assumed to behave incompressibly (Budday et al., 2017a; Feng et al., 2017). This enables the use of available closed form solutions for simple load cases. In this work, we fitted the compressible one-term Ogden model with different levels of compressibility, i.e., *v* = {0.45,0.49}. A slightly compressible formulation circumvents numerical problems associated with incompressibility.

#### Boundary conditions: “glued” vs. “slipping”

In our experiments, the specimens have to be glued to the specimen holders to enable tensile testing. Therefore, the bottom and upper surface are fixed, which results in an inhomogeneous deformation state. To check the influence of the glued in comparison to ideal slipping boundary conditions we ran finite element simulations for both scenarios. In the case of ’glued’, non-slipping boundary conditions, all nodes on the top and bottom surface are held fixed. For the slipping boundary conditions, nodes are only fixated in all directions along the centerline of the cylinder and can freely move in the plane of the bottom and top surface, where only the axial direction is fixed. Furthermore, they are fixed in the normal directions of the planes orthogonal to the cylinder axis to prevent rigid body motion.

### Data preprocessing: The hyperelastic response

As we limit ourselves to a hyperelastic material model (neglecting poro- and viscoelastic effects), we need to extract the hyperelastic response from the experimental data, which shows a considerable hysteresis (see Figure 3). The (theoretically infinitely) high or low strain rates that would be needed to obtain a purely hyperelastic response are not feasible in experiments. Figure 3 gives a short overview over the data after the processing steps from the raw data as it is obtained from the rheometer to the final processed data. We adopt here the assumption that the averaged loading and unloading curves approximate the hyperelastic response (Budday et al., 2017a). The initial moving average and lowpass filtering helps to filter out the high frequency noises in the measured signal. A resampling along the displacement axes enables the averaging of loading and unloading to finally obtain our desired results. The number of points is reduced to 60 points per mode so that the computational costs for the simulation are lowered, while the characteristic shape of the curve is preserved. We use either the data from the first or the third loading cycle to represent the un- and preconditioned material response, respectively. Together with the two used Poisson’s ratios *v* = {0.45,0.49} this leads to a total of four parameter sets for each specimen.

**Figure 3.**
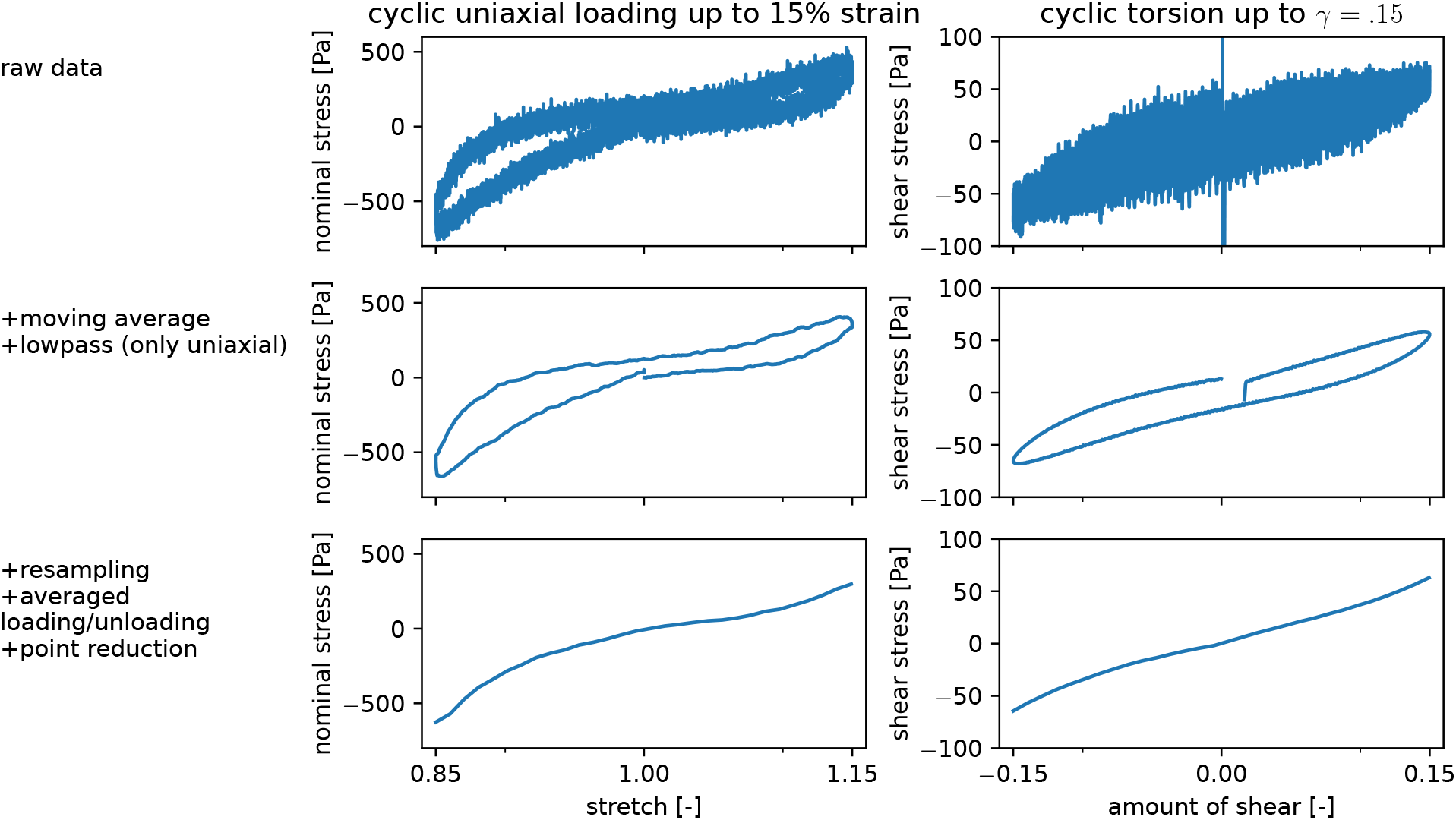
Preprocessing steps from raw data (top row) to the averaged unloading and loading curves representing the hyperelastic response (bottom row), exemplary shown for data from the first cycle of compression-tension, and torsional shear for one representative specimen from the putamen (Pu).

### Inverse parameter identification

To characterize the mechanical behavior of the tested human brain specimens, we inversely determine material parameters for the modified one-term Ogden model, as summarized in Figure 4. In the generalized problem, the model *G*(**m**) depends on the parameters **m** and produces the results *d*,

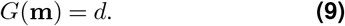

**Figure 4.**
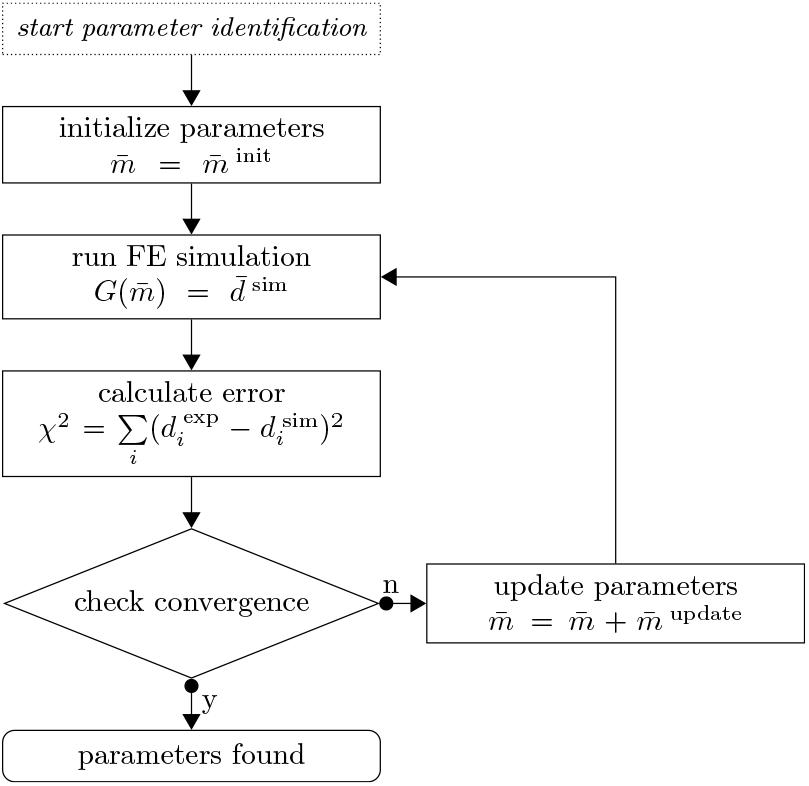
Parameter identification scheme. The set of material parameters 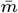 is updated in each iteration by the used optimization algorithm until a convergence criterion is met.

In the real setup, our measured experimental output differs from the output produced using the ’true’ parameter set **m**^true^ by the error *η*

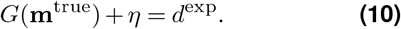

Our goal is now to find the optimal parameter set **m**\* that best reproduces the output of our experiments *d*^exp^. We assume the error to be normally distributed 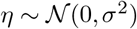 with the mean at 0 and the standard deviation *σ*. The solution of the identification problem will then approximate **m**^true^ when an L2-Norm is used to measure the error between the simulated values and the experimental data following the maximum likelihood principle (Seber et al., 2005). We measure the goodness of fit by the normalized squared error of experimental and model output values

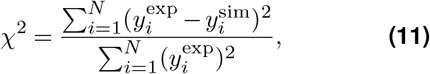

where *y*^sim^ and *y*^exp^ denote the simulation output and the corresponding value measured during the experiment, respectively. The approach of normalizing the squared error was taken from Gavrus et al., 1996 and helps to tackle numerical problems like vanishing gradients that may arise for low values of *χ*^2^.

The parameter identification is implemented in Python and coupled with the finite element simulation using the .prm file format of the deal.ii library. We use the implementations of optimization routines in the SciPy Python-module (Virtanen et al., 2020) and adapt them to improve computational efficiency by the parallel evaluation of finite element simulations. We note that we have initially used the gradient-free Nelder-Mead algorithm (for brain 1,2 and 3 using *scipy.optimize.minimize* with the options *method*=’Nelder-Mead’, *xatol* = 10^-3^ and *fatol* = 10^-4^) described by Nelder et al., 1965, as we had problems with vanishing gradients which did not allow us to use gradient-based algorithms. This algorithm has already been used successfully for the parameter identification of brain tissue (Prevost et al., 2011; Hosseini-Farid et al., 2019). After the introduction of the aforementioned normalization factor 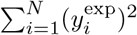 in *χ*^2^, those problems were overcome. Consequently, we switched to the trust region reflective algorithm, formulated by Branch et al., 1999, which can handle constraints and appears to be faster, as less iterations were needed to fulfill the set convergence criterion. Our adaptation of the algorithm is mainly the switch to an eager evaluation of function values. This means that a new simulation run will be started as a new asynchronous thread as soon as we know that this value will be needed although the output might not be processed at the same place in the code. When the output is then actually needed in a later step, the return of the asynchronous process is awaited. Furthermore, simulation outputs are cached to prevent unnecessary simulation runs. The gradient is calculated using a finite forward difference scheme. Importantly, a preliminary global optimality study, where the optimization algorithm was started with varying initial parameters, showed no problems of multiple local minima (Supplementary Figure S14).

In a first step, we apply the inverse parameter identification scheme to the data from each of the 182 human brain specimens individually. Statistical methods allow us to quantify significant differences between parameter sets and their subgroups (regions, brains). We use the Python module pandas (McKinney, 2010) for data analysis and the modules seaborn (Waskom, 2021) and matplotlib (Hunter, 2007) for visualizations. For each of the specimens, we consider the data from three loading modes simultaneously, i.e., compression and tension loading up to 15% nominal strain as well as torsional shear loading up to an amount of shear of *γ* = {0.15,0.3}. We do not introduce any explicit weighing mechanisms into the cost function *χ*^2^. As all three modes contribute with the same number of data points to the residual vector, in fact, shear loading is implicitely weighed with a ratio of 2 : 1 compared to the compression and tension loadings. We achieved good results using this approach that can be justified by the maximum compression stresses reaching a multiple in magnitude compared to the shear stresses. We note that the weighing of the loading modes has an important influence on the identification of material parameters and can be adjusted to the intended purpose. As we aim to identify a universal parameter set without having a specific application in mind, we did not use any explicit weighing. This could be reasonable, however, if a specific application has a known dominant loading mode.

### Statistical analyses

We first check if our data is normally distributed as parametric statistical tests that rely on the assumption that the tested data is drawn from a normal distribution are favored for their higher statistical power (King et al., 2019). To this end, we apply a Shapiro-Wilk test using the implementation provided by the SciPy Python module (Virtanen et al., 2020) to the material parameters of the Ogden model *α* and *μ* as well as the error of the fit in terms of the root mean square error (RMSE). As this analysis lead us to reject the assumption of a normal distribution for the material parameters, we will introduce the used non-parameteric tests in the following. The Kruskal-Wallis H-test is employed as a nonparametric equivalent of an ANOVA to determine if the means of the groups defined as the anatomical regions, governing regions and brains are significantly different. The test is run for both material parameters *α* and *μ*, for each group and for all four parameter sets separately as they are not independent. To identify, which governing regions should be treated as different regarding their hyperelastic behavior, we conduct pairwise post hoc tests in terms of Mann-Whitney-U tests. Additionally, the same test is used to quantify pairwise differences between the parameters of the tested brains. We use the implementation of the method in the Python module scikit_posthocs (Terpilowski, 2019), where we use the Holm-Bonferroni method to control the family-wise error rate. By calculating the pairwise differences between the four parameter sets we are able to quantify the influence of preconditioning and the two Poisson’s ratios on the material parameters. To this end, we use a Wilcoxon signed-rank test. The returned *p*-value is an estimate on the probability that these observations come from a distribution with a mean value different from 0. We use the *wilcoxon* function in the SciPy Python module (Virtanen et al., 2020). We consider a *p*-value lower than 0.05 to be significant for all conducted tests.

## Results

### The effect of inhomogeneous deformation states during testing

Figure 5 demonstrates the significant influence of the boundary conditions during compression and tension loadings, where the absolute nominal stresses for glued boundary conditions are twice as high as for slipping boundary conditions – corresponding to homogeneous deformation states. The maximum local stresses visualized in Figure 5c are an order of magnitude higher than in Figure 5b, which can be attributed to the singularity at the edge of the fixed surface. We note that we only present results for compression and tension loadings, as the torsional loading will not be influenced by the two types of boundary conditions. These results highlight the importance of performing an inverse parameter identification scheme, as presented in the following, instead of using closed form analytical solutions based on the assumption of homogeneous deformation states.

**Figure 5.**
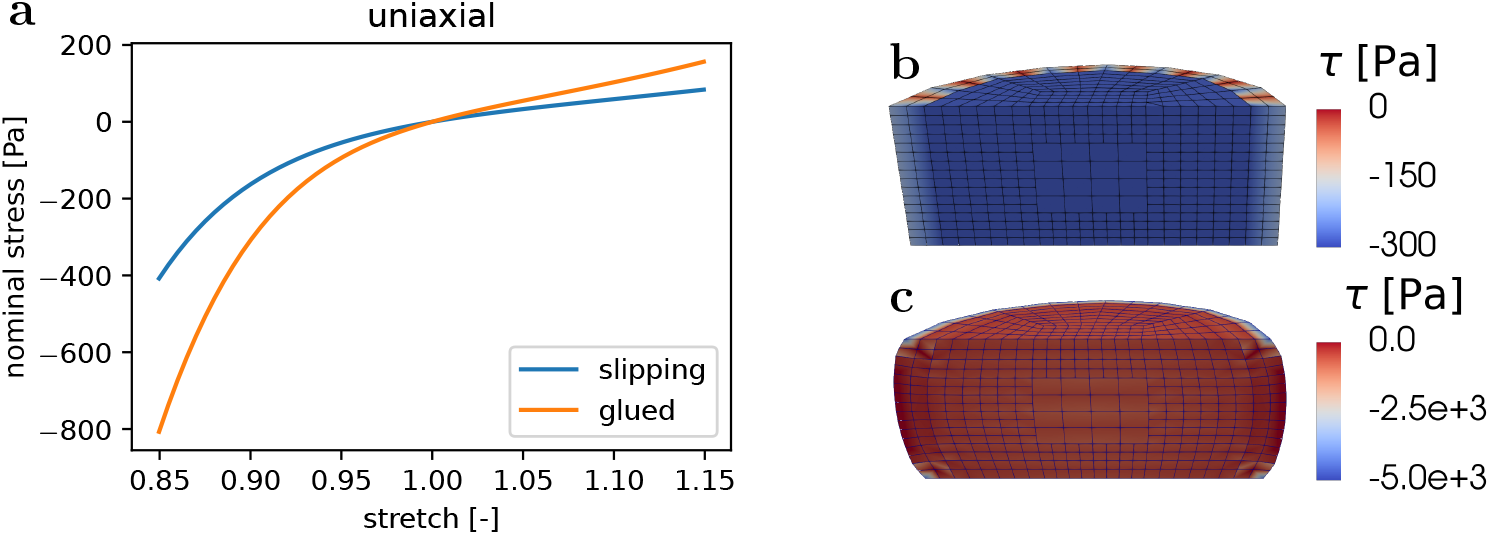
The direct comparison shows that ’glued’, non-slipping boundary conditions lead to a stiffer response than ’slipping’ boundary conditions. (**a**) Simulated stress-stretch results for the ’glued’ and ’slipping’ boundary conditions. (**b**) ’Slipping’ boundary conditions lead to a homogeneous deformation state. (**c**) ’Glued’ boundary conditions lead to an inhomogeneous deformation state.

### Performance of parameter identification scheme and general trends

Figure 6 shows an exemplary model fit for one of the specimens compared to the preprocessed experimental data. The model well captures the experimental response, including the pronounced compression-tension asymmetry. Differences between the four parameter sets clearly appear in the scatter plots in Figure 7, where the parameters of each specimen are represented by a dot. While the nonlinearity parameter *α* shows no clear trend, the change in the shear modulus *μ* is already visible in this representation. For both values of the Poisson’s ratio *v*, we observe lower shear moduli when the model is fitted to preconditioned data. Furthermore, the higher Poisson’s ratio leads to a more incompressible and therefore stiffer behavior of the model, which is countered by lower shear moduli in the parameter sets for a Poisson’s ratio of 0.49.

**Figure 6.**
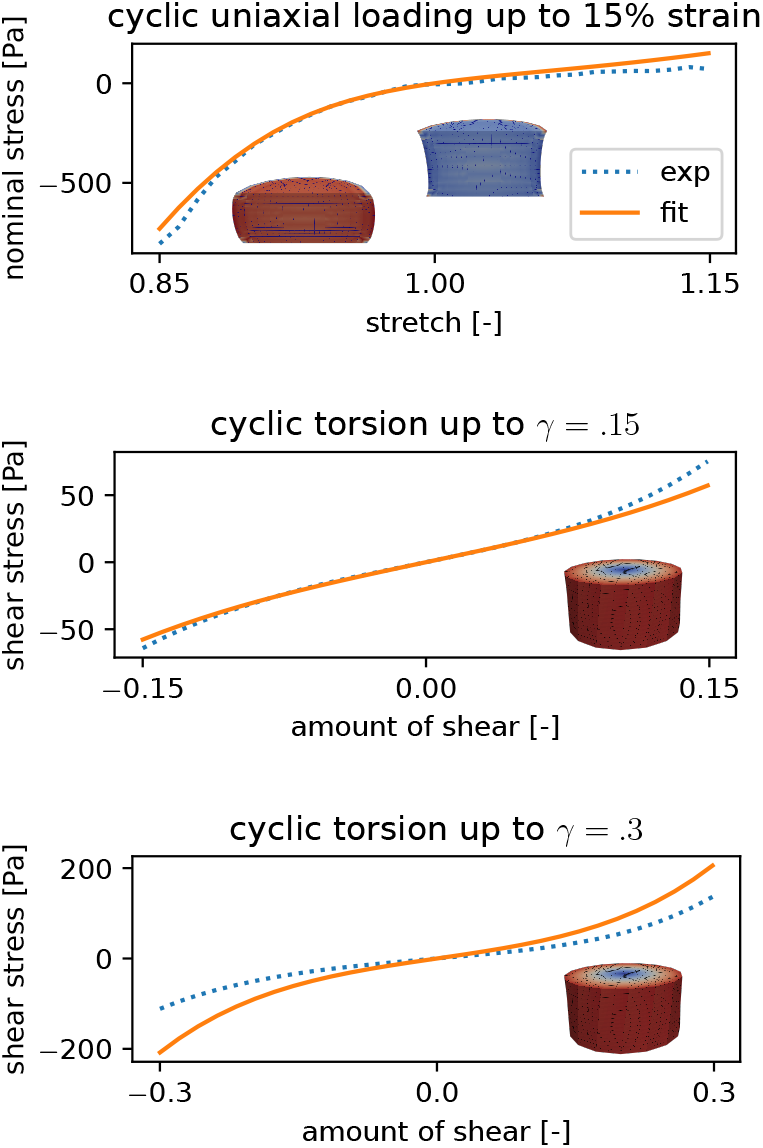
Exemplary fitting results for one specimen from the midbrain, where the preconditioned response was fitted using a Poisson’s ratio of 0.45.

**Figure 7.**
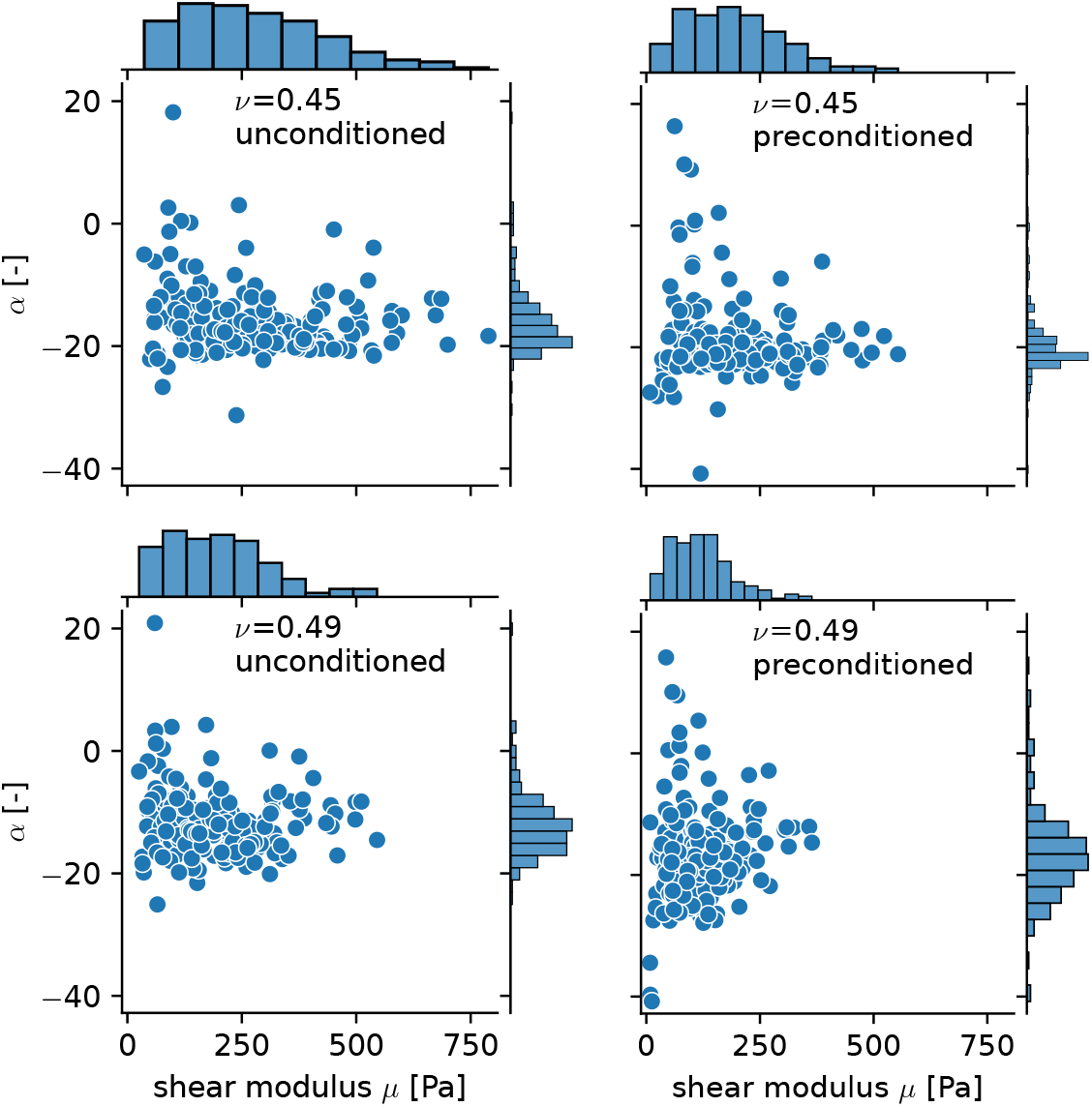
Overview of the obtained four sets of material parameters.

### Regional dependency

Figure 8 (top) shows the material parameters that were fitted to the unconditioned data (first cycle) using a Poisson’s ratio of 0.45 averaged over the 19 anatomical regions introduced in Table 3. We only plot this data set here as the other three cases (Poisson’s ratio 0.49 and preconditioned) show the same qualitative behavior (see also Supplementary Figures S1 and S2). The (anatomical) region-wise averaged values of the model parameters can be found in Supplementary Table S2. From the overlapping bars indicating the standard deviation in Figure 8 (top), it is clear that this partitioning is too detailed to identify regions with distinct hyperelastic properties. Therefore, we subsequently group regions with similar parameters as well as comparable microstructures and location into what we define as nine ’governing’ regions. Table 5 lists the used abbreviations as well as the assignment of regions to governing regions. The distribution of governing regions over the tested brains is shown in Figure 9. The distribution over the tested brains is similar besides the Amygdala (Am) region, for which the data set contains only samples from the brains 3, 4, 5, and 7. An equal distribution is important for the conducted statistical analyses. The resulting distribution of parameters in Figure 8 (bottom) shows a slight improvement in terms of reducing overlapping, but especially the samples from the midbrain (M), the brainstem (BS) and the basal ganglia (BG) governing regions are still close in the parameter space and show overlapping standard deviations. The corresponding averaged material parameter values can be found in Supplementary Table S1.

**Table 5.** Governing regions and their abbreviations with corresponding anatomical regions.

| Abbreviation | Full name | Assigned regions |
| --- | --- | --- |
| Am | Amygdala | Am |
| BG | Basal ganglia | Pa,Pu,NC |
| BS | Brain stem | Me,P |
| C | Cortex | MC,VC,CI,FC,TL |
| CB | Cerebellum | cWM,cN |
| CC | Corpus callosum | CC |
| CR | Corona radiata | CR,WM |
| Hi | Hippocampus | Hi |
| M | Midbrain | M,Th |

**Figure 8.**
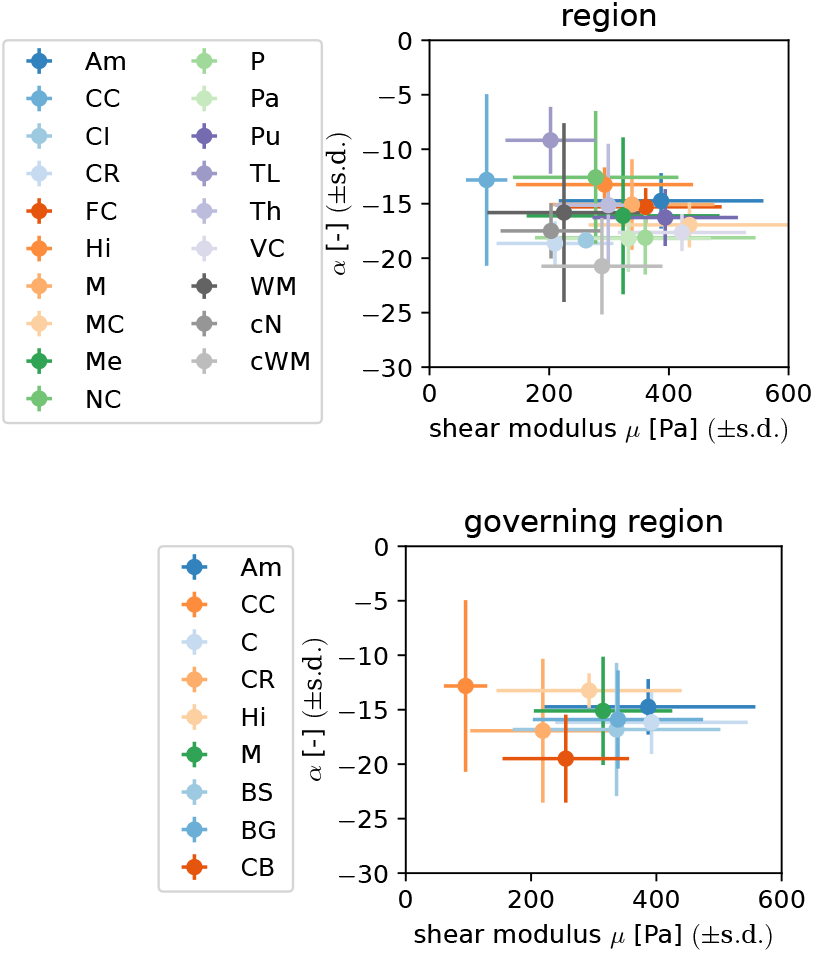
Material parameters for the anatomical and defined governing regions, obtained by fitting the unconditioned data using a Poisson’s ratio of 0.45.

**Figure 9.**
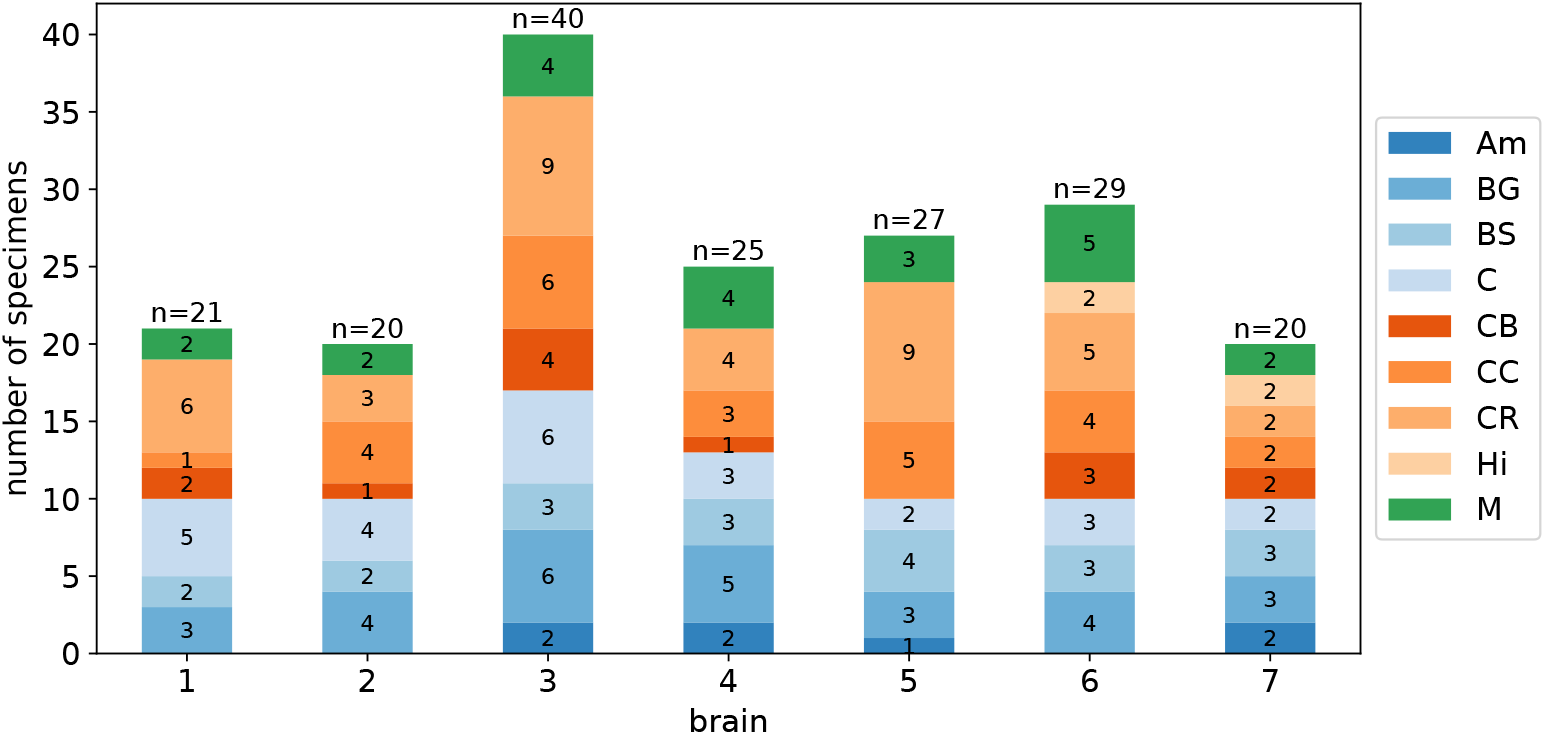
Distribution of specimens from governing regions for the individual brains.

**Figure 10.**
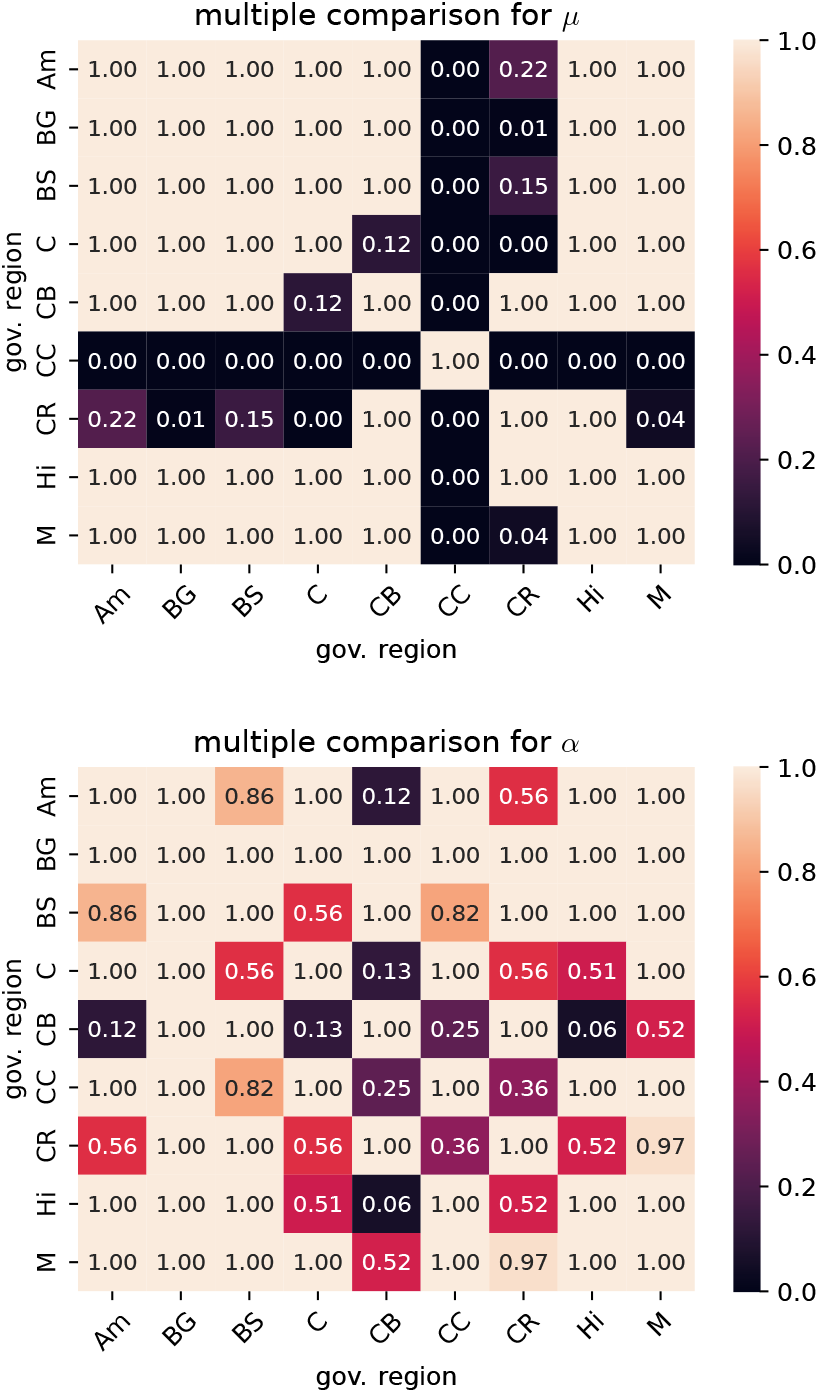
*p*-values from pairwise post hoc Mann-Whitney-U tests comparing material parameters for different governing regions for a Poisson’s ratio of 0.45 fitted to the unconditioned data.

Table 6 and Table 7 list the H-statistics of the Kruskal-Wallis H-test as well as the corresponding *p*-values when the detailed anatomical and governing regions were used as grouping variable, respectively. All *p*-values are below the significance level. While the *p*-values for the governing regions are lower than the corresponding values for the detailed regions, it is difficult to directly interpret this qualitative difference, as the results of the statistical test are influenced by the number of groups as well as the number of observations inside each group. To identify, which governing regions should be treated as different regarding their hyperelastic behavior, we subsequently conduct pairwise post hoc tests in terms of Mann-Whitney-U tests. Figure 10 shows the corresponding results as matrix plots for unconditioned data and a Poisson’s ratio of 0.45. Specimens from the corpus callosum (CC) governing region show significantly different shear moduli in all comparisons. Shear moduli for the corona radiata (CR) are significantly different compared to the basal ganglia (BG), cortex (C), corpus callosum (CC) and midbrain (M), but not to the other governing regions. The shear moduli of the remaining seven governing regions do not appear to be significantly different in the conducted statistical tests. Multiple comparison tests for the nonlinearity parameter *α* did not show any of the governing regions to have significantly different values. The other three parameter sets, for which the results of the multiple comparison tests are presented in Supplementary Figures S5 and S6, show the same qualitative behavior with only small variations in the number of significant comparisons. One exception are two significant comparisons for *α* for the parameter set of the preconditioned data and a Poisson’s ratio of 0.45, which do not appear for any of the other parameter sets.

**Table 6.** Results for the Kruskal-Wallis H-test with the anatomical region as independent variable

| $\nu$ | preconditioned | $\alpha$ | | $\mu$ | |
| --- | --- | --- | --- | --- | --- |
|  |  | H | p | H | p |
| 0.45 | preconditioned | 3.27e+01 | 1.83e-02 | 9.20e+01 | 6.31e-12 |
| 0.45 | unconditioned | 3.92e+01 | 2.67e-03 | 8.72e+01 | 4.61e-11 |
| 0.49 | preconditioned | 3.57e+01 | 7.67e-03 | 1.05e+02 | 2.38e-14 |
| 0.49 | unconditioned | 4.27e+01 | 8.79e-04 | 9.20e+01 | 6.25e-12 |

**Table 7.** Results for the Kruskal-Wallis H-test with the governing region as independent variable

| $\nu$ | preconditioned | $\alpha$ | | $\mu$ | |
| --- | --- | --- | --- | --- | --- |
|  |  | H | p | H | p |
| 0.45 | preconditioned | 1.92e+01 | 1.38e-02 | 8.07e+01 | 3.61e-14 |
| 0.45 | unconditioned | 2.48e+01 | 1.65e-03 | 7.85e+01 | 9.78e-14 |
| 0.49 | preconditioned | 2.35e+01 | 2.73e-03 | 9.53e+01 | 3.85e-17 |
| 0.49 | unconditioned | 2.86e+01 | 3.67e-04 | 8.30e+01 | 1.21e-14 |

### Inter-individual variation

When aiming to choose material parameters for human brain models, an important question is whether the parameters can be generalized or need to be patientspecific. Figure 11 shows an overview of the parameters identified for the seven different brains. Again, we only discuss the parameter set of the unconditioned response using a Poissons’s ratio of 0.45 in detail, while the results of the remaining parameters can be found in Supplementary Figure S7. The shear moduli range from 185Pa for brain 3 to 365Pa for brain 4 and the nonlinearity parameter *α* ranges from −18 for brain 4 to −13 for brain 7. The brain-specific parameters are not spread evenly throughout the parameter space with brains 1,4,5 and 6 having relatively high shear moduli and lying close together, while brain 3 and 7 show lower shear moduli. Brain 2 is somewhere in between these two groups. The *p*-values returned by a Kruskal-Wallis H-Test with the individual brains as independent variable in Table 8 are below the significance level for both parameters *α* and *μ* and for all four parameter sets. The results of pairwise Mann-Whitney-U tests in Figure 12 show that mainly the shear moduli for brain 3 significantly differed from all other brains, excluding 2 and 7. Another significant comparison is found between 4 and 7. For the nonlinearity parameter *α*, most significant differences are found for brain 7. Furthermore, the comparison of brain 4 and 6 is reported as significant. The results for the other three parameter sets can be found in Supplementary Figures S8 and S9.

**Figure 11.**
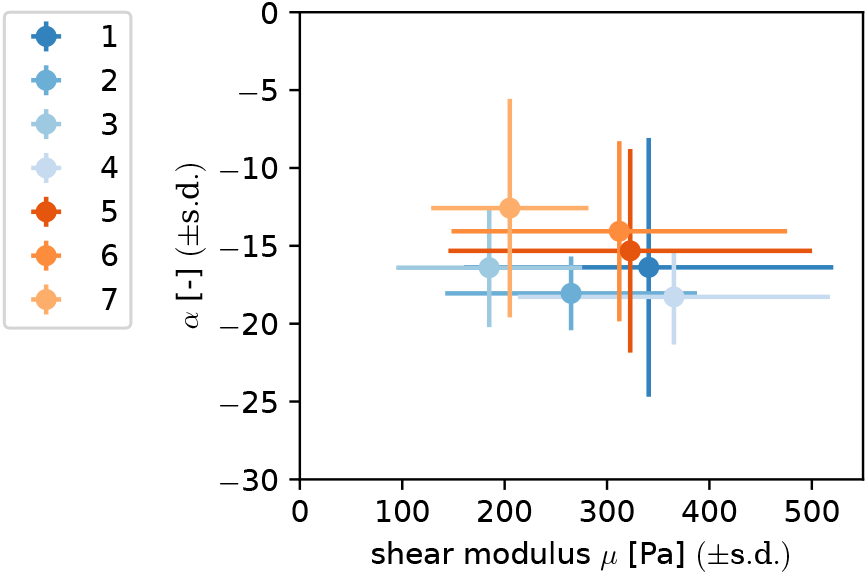
Parameter values and their standard deviation after averaging over different individual brains for the unconditioned response and a Poissons’s ratio of 0.45.

**Figure 12.**
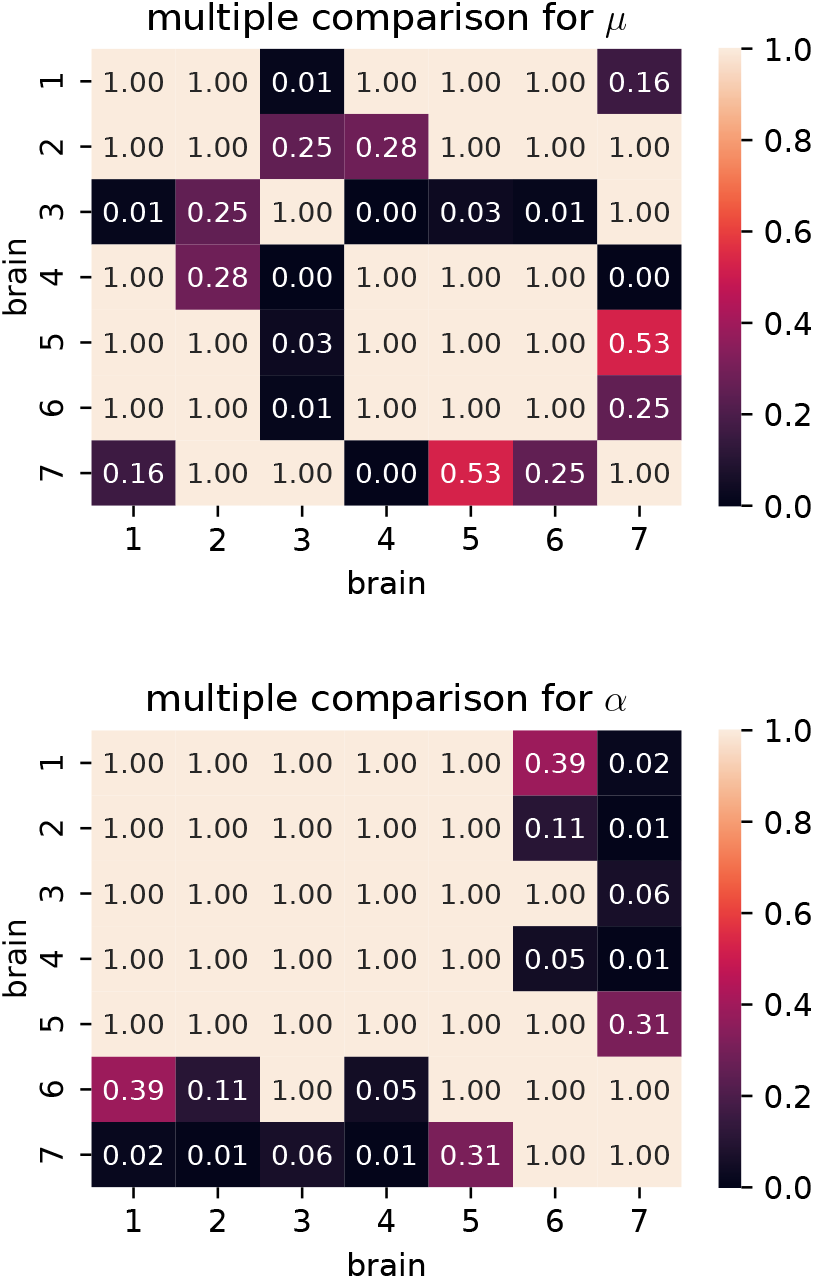
*p*-values from pairwise post hoc Mann-Whitney-U tests comparing material parameters from different brains for a Poisson’s ratio of 0.45 fitted to the unconditioned data.

**Table 8.** Results for the Kruskal-Wallis H-test with the brain as independent variable

| $\nu$ | preconditioned | $\alpha$ | | $\mu$ | |
| --- | --- | --- | --- | --- | --- |
|  |  | H | p | H | p |
| 0.45 | preconditioned | 1.95e+01 | 3.41e-03 | 3.65e+01 | 2.16e-06 |
| 0.45 | unconditioned | 2.48e+01 | 3.65e-04 | 3.31e+01 | 1.02e-05 |
| 0.49 | preconditioned | 1.59e+01 | 1.43e-02 | 2.42e+01 | 4.78e-04 |
| 0.49 | unconditioned | 3.16e+01 | 1.93e-05 | 3.10e+01 | 2.57e-05 |

### Effect of compressibility

We fitted the experimental results using different compressibilities, quantified through the Poisson’s ratio *v*, which we relate to the shear modulus *μ* of the modified one-term Ogden model by Equation 8 known from the linear regime. We use the Poisson’s ratios 0.45 and 0.49 to enable the comparison with similar studies but would like to note that this does not directly conform to the definition in terms of the ratio of transverse and axial strain. With these two parameter sets, we are able to evaluate the effect of a prefixed Poisson’s ratio on obtained material parameters for each specimen and to see whether one of those leads to lower RMSE and thus better quality of the fit. In the following, we only focus on the unconditioned data as the found relations also hold qualitatively for the preconditioned data. The complete results can be found in Supplementary Figure S10. Figure 13 shows the histograms of the pairwise differences for the unconditioned data with their median. The boxplot of the RMSE values puts the calculated differences into perspective. RMSE values for unconditioned data have a mean of 48Pa for a Poisson’s ratio of 0.45 and 34Pa for 0.49. The *p*-values of a Wilcoxon signed-rank test in Table 9 indicate all observed differences as significant. The median of the difference is −71Pa for the shear modulus, 4 for the nonlinearity parameter *α* and −12Pa for the RMSE. A negative difference corresponds to a higher value for a Poisson’s ratio of 0.45.

**Table 9.** Results of the Wilcoxon signed-rank test comparing the datasets for the Poisson’s ratios *v* = {0.45,0.49}.

| preconditioned | $H_0 := \Delta\alpha = 0$ | | $H_0 := \Delta\mu = 0$ | | $H_0 := \Delta RMSE = 0$ | |
| --- | --- | --- | --- | --- | --- | --- |
|  | W | p | W | p | W | p |
| preconditioned | 2.88e+03 | 2.03e-14 | 1.13e+02 | 8.24e-31 | 2.61e+03 | 1.00e-15 |
| unconditioned | 7.30e+01 | 4.28e-31 | 1.62e+02 | 1.83e-30 | 2.58e+03 | 6.56e-16 |

**Table 10.**
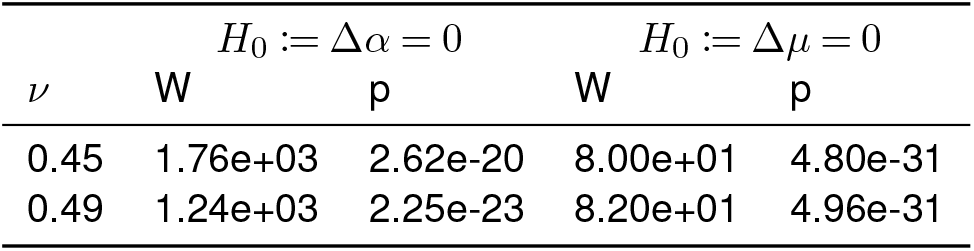
Results of the Wilcoxon signed-rank tests comparing the datasets for un- and preconditioned data.

**Figure 13.**
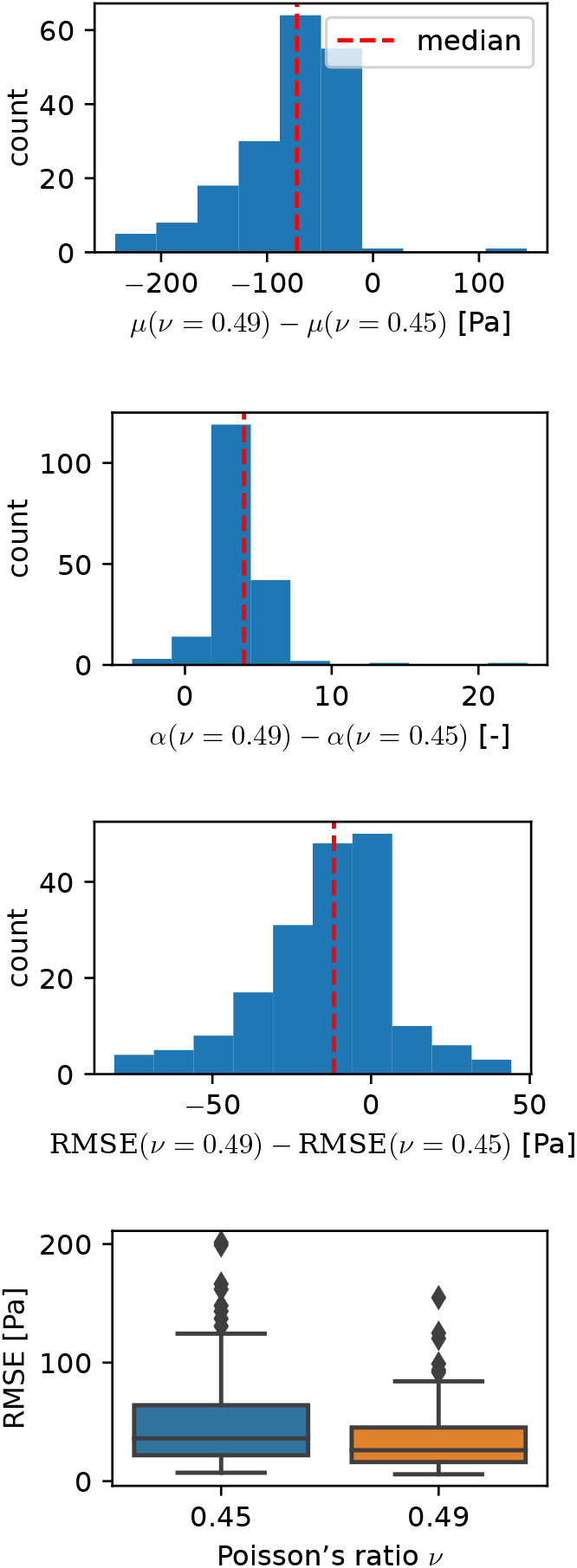
Pairwise difference for the parameters *μ* and *α* and the RMSE between samples fitted with a Poisson’s ratio of 0.45 and 0.49. Results are shown for the unconditioned data set. The boxplot (bottom right) visualizes the distribution of RMSE values.

### Preconditioned versus unconditioned material parameters

We obtained separate parameter sets for the unconditioned and preconditioned material responses by fitting the first and third loading cycle of all loading modes, respectively. Figure 14 shows the histogram of the pairwise differences between material parameters for a Poisson’s ratio of 0.45. The results for a Poisson’s ratio of 0.49 show the same qualitative trends and are visualized in Supplementary Figure S11. Table 10 shows the results of a Wilcoxon signed-rank test, indicating all observed differences as significant. The differences for the shear moduli and nonlinearity parameter *α* have both negative medians with Δ*μ* = −74Pa and Δ*α* = −2.7, indicating lower values for the preconditioned data.

**Figure 14.**
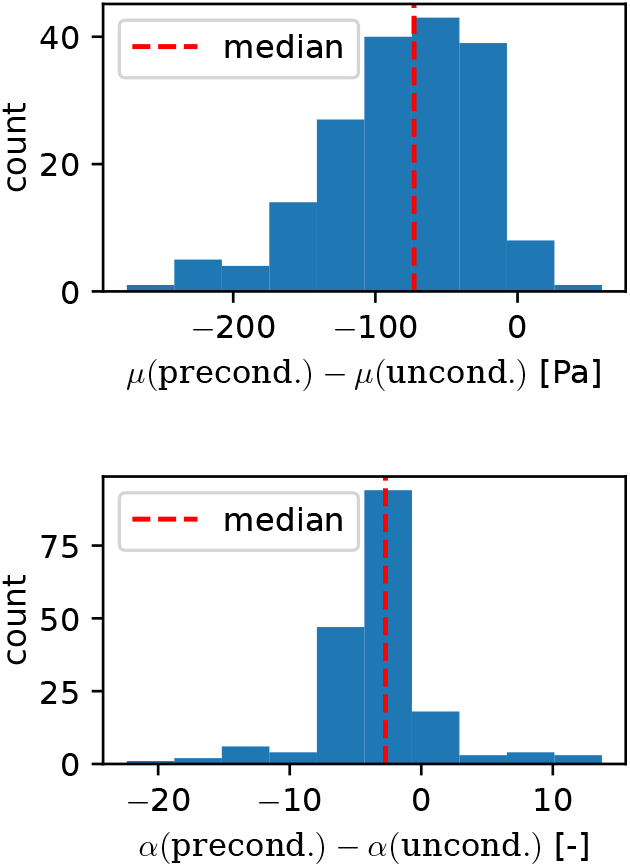
Pairwise difference in the material parameters between samples fitted to the un- and preconditioned data for a Poisson’s ratio of 0.45.

### Averaging material parameters versus averaging experimental data

While we fitted the experimental output of every single specimen separately in a first step to perform statistical analyses, this approach is not accurate when aiming to provide material parameters representing the averaged response of each of the defined regions. Therefore, in a second step, we applied the parameter identification scheme after averaging the experimental data over the governing regions. Figure 15 compares the parameters from both approaches for the unconditioned data using a Poisson’s ratio of 0.45. The maximum absolute relative differences are 13% for *μ* and 33% for *α*. The differences for the other three data sets can be found in Supplementary Figures S12 and S13. Over all data sets, the maximum absolute relative difference for *μ* is 32%. All parameters obtained from fitting the averaged experimental results were within the standard deviation of averaging the individual specimen-specific parameters. Interestingly, the averaged data resulted in consistently higher absolute values of the nonlinearity parameter *α*, but lower shear moduli *μ*. The parameter values obtained when fitting the averaged experimental results in the governing regions are summarized in Table 11.

**Figure 15.**
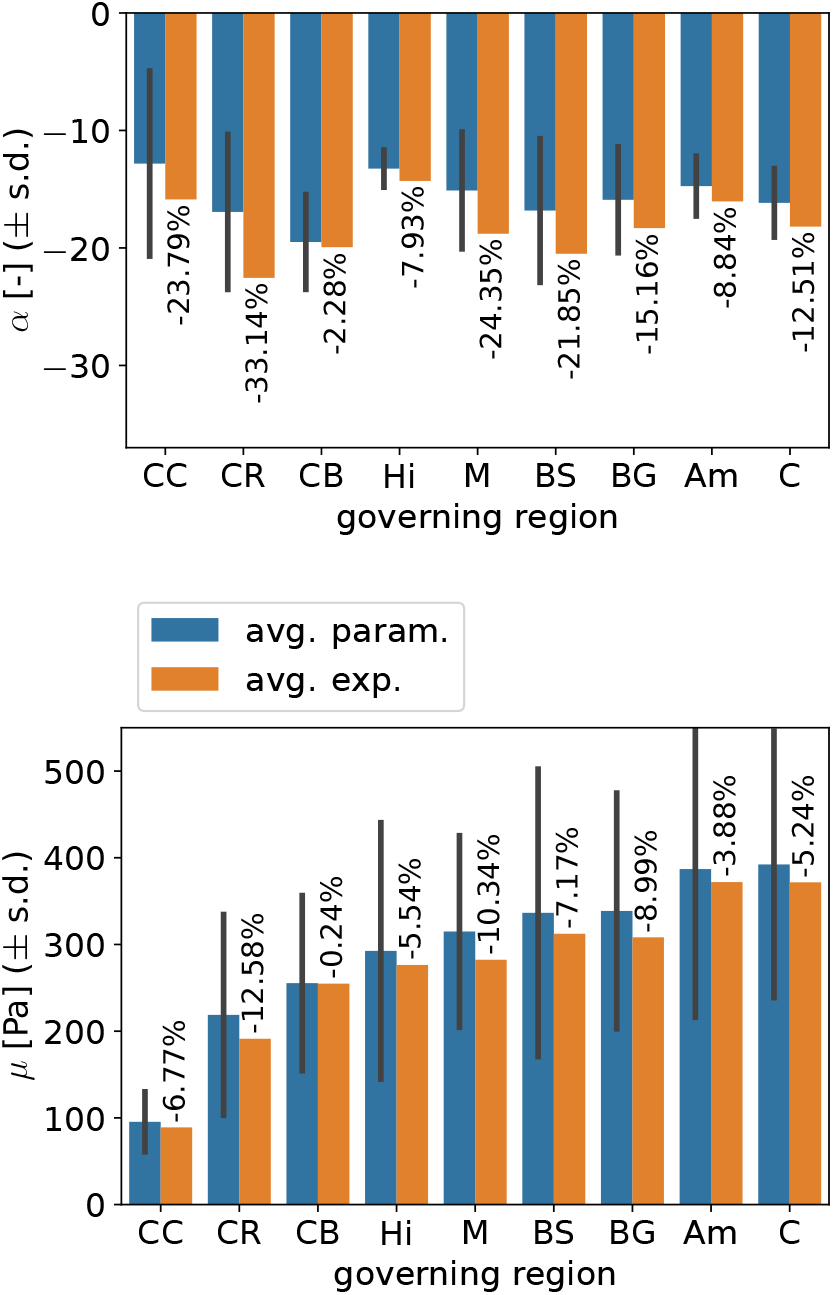
Comparison between the averaged material parameters obtained from fitting the experimental data of each specimen separately and those obtained from fitting the averaged experimental response. Results are shown for the parameter set fitted to the unconditioned response using a Poisson’s ratio of 0.45.

**Table 11.** Material parameters fitted to the experimental response averaged over governing regions.

| gov. region | preconditioned $\nu = 45$ | | preconditioned $\nu = 49$ | | unconditioned $\nu = 45$ | | unconditioned $\nu = 49$ | |
| --- | --- | --- | --- | --- | --- | --- | --- | --- |
| | $\alpha$ | $\mu$ | $\alpha$ | $\mu$ | $\alpha$ | $\mu$ | $\alpha$ | $\mu$ |
| Am | -20.70 | 234.92 | -13.96 | 146.04 | -16.04 | 371.94 | -10.72 | 283.86 |
| BG | -21.41 | 218.73 | -15.80 | 126.24 | -18.31 | 308.21 | -12.21 | 236.32 |
| BS | -22.70 | 209.41 | -25.25 | 79.81 | -20.49 | 312.33 | -14.98 | 227.35 |
| C | -21.10 | 258.77 | -17.79 | 136.07 | -18.18 | 371.68 | -12.76 | 279.48 |
| CB | -22.40 | 171.48 | -20.72 | 79.49 | -19.94 | 254.77 | -14.11 | 187.94 |
| CC | -19.04 | 51.80 | -14.90 | 29.48 | -15.87 | 88.97 | -11.19 | 66.68 |
| CR | -24.72 | 132.34 | -21.60 | 61.47 | -22.54 | 191.24 | -15.18 | 142.67 |
| Hi | -18.99 | 163.29 | -13.37 | 99.49 | -14.30 | 276.37 | -9.66 | 209.32 |
| M | -22.63 | 182.80 | -17.83 | 98.31 | -18.79 | 282.36 | -12.48 | 216.84 |

## Discussion

In this work, we have performed multi-modal large-strain mechanical testing of human brain tissue and have implemented an inverse parameter identification scheme utilizing a finite element implementation of the modified one-term Ogden model to (i) identify mechanically distinct human brain regions, and (ii) provide the corresponding hyperelastic material parameters for future finite element simulations. Each specimen was tested under multiple consecutive loading modes, namely cyclic compression and tension loading as well as cyclic torsional shear, which required to glue the specimens to the specimen holders during testing. Therefore, we have carefully analyzed the validity of assuming a homogeneous deformation state, as often done in the literature. In addition, we have investigated the effect of predefining different Poisson’s ratios, and have provided separate parameter sets for the unconditioned and preconditioned material responses, respectively.

### The importance of an inverse parameter identification scheme

It is common to characterize the mechanical properties of materials by testing them under simple loading modes, where it is possible to obtain analytical solutions based on certain assumptions. If the deformation gradient **F** is assumed to be homogeneous (therefore describing an affine deformation) for a hyperelastic material, the stress will also be homogeneous and can be directly obtained from a constitutive relation by using Equation 5. Whenever possible, the application of such closed form solutions is favorable over computationally expensive approaches like the finite element method. However, it needs to be checked how well the assumption of homogeneous deformation approximates the real conditions during the experiment. The model outputs in Figure 5 for slipping and non-slipping boundary conditions show a strong influence with approximately two times higher nominal stresses for the case when specimens are glued on the top and bottom surfaces. These results emphasize the importance of using an inverse parameter identification scheme based on a model that can accurately capture the boundary conditions during testing. Similar results were obtained by other researchers (Miller, 2005; Voyiadjis et al., 2018; Felfelian et al., 2019; Budday et al., 2019).

### Mechanically distinct brain regions

From the initially considered 19 anatomically different brain regions, we were able to assign nine governing regions grouping specimens that were extracted from anatomical regions with comparable microstructures as well as similar mechanical parameters. Our results indicate that from these governing regions at least the corpus callosum and the corona radiata should be modeled as distinct regions in mechanical models of the human brain. The corpus callosum yields the lowest shear moduli for all four datasets. While we obtain the cortex (C) as region with the highest shear moduli for a Poisson’s ratio of 0.45, the Amygdala (Am) is the stiffest region for 0.49. Interestingly, as the significant differences were only found for the shear modulus *μ*, the use of a constant value for *α* over all regions may be justified when using a one-term Ogden model. The observed trends generally agree with previous studies investigating a limited number of brain regions. For instance, Budday et al., 2017a showed significant differences between the corpus callosum and both the basal ganglia as well as the cortex from compression, tension, and shear tests. Interestingly, differences between the corona radiata and corpus callosum were also reported as significant by pairwise tests but could not be confirmed by multiple comparisons. Another study identified viscoelastic parameters from indentation experiments distinguishing 12 regions with six of them being subregions of the cortex (Menichetti et al., 2020). Multiple comparison tests were conducted, where the instantaneous shear modulus *μ*_0_ for every region was significantly different from at least three other regions. Most significant comparisons were found for the corona radiata and the cerebellum with seven, whereas only three were found for the corpus callosum. The reported difference in the region-specific mechanical behavior of brain tissue when comparing indentation with compression, tension and shear loadings could explain the differences in the identified mechanically distinct regions. Still, the corona radiata was identified as a mechanically distinct region in both cases.

### Region-specific parameters

After we have identified mechanically distinct regions by means of statistical analyses of the parameters obtained for each specimen individually, we subsequently provide (governing) region-specific hyperelastic parameters based on the averaged data of all specimens within the respective region, as reported in Table 11. The qualitative regional trends are comparable to our previous results in (Budday et al., 2017a), where we had reported one-term Ogden parameters fitted to compression, tension and shear data for the corpus callosum, basal ganglia, cortex and the corona radiata. Still, the shear moduli were substantially higher ranging from 4.1 fold for the basal ganglia to 9.3 fold for the corpus callosum in comparison to our dataset for a Poisson’s ratio of 0.49 and preconditioned data. This can be attributed to the different modeling approaches as we had used an incompressible analytical implementation assuming slipping boundary conditions. Another study (Moran et al., 2014) reported parameters for a two-term Ogden model fitted to tension, compression and shear tests of human brain tissue, where they distinguished the corona radiata as well as gray and white matter. The shear moduli, calculated as *μ* = *μ*_1_ + *μ*_2_, were higher for gray matter than for white matter, which can be considered to be in agreement with our study. We identified the lowest shear moduli for the white matter regions corpus callosum and corona radiata, while the highest values were found for the gray matter regions cortex and amygdala. Their reported values for the corona radiata, which is already contained in the white matter data but again introduced as a separate region, are the highest. All values reach more than three fold of our values.

### Influence of compressibility

By fitting the experimental data with the two different fixed Poisson’s ratios 0.45 and 0.49, we were able to firstly investigate whether one of them fitted the data better and, secondly, how the chosen Poisson’s ratio affects the identified remaining free model parameters. We note that the Poisson’s ratio is here used to relate the shear modulus *μ* to the bulk modulus *κ* in terms of Equation 8, thus actually denoting the initial Poisson’s ratio in the finite strain regime.

A potential influence on the quality of the fitting results is quantified in terms of differences in the root mean squared error (RMSE) between experimental data and model output for each specimen and all loading modes. Although the reported negative median indicates a slightly better fit for a Poisson’s ratio of 0.49, the long tails in the histogram of the differences in Figure 13 and Supplementary Figure S10 as well as the large spread in the obtained RMSE values, visualized by the boxplots, do not support a systematic dependency. This shows that the available experimental data does not contain enough information to characterize the volumetric part of the constitutive model. A parameter study on a neo-Hookean model replicating indentation experiments varied the Poisson’s ratio between 0.452 – 0.49995 (MacManus et al., 2018). The increase in maximum indentation force by 6% was below the coefficient of variation of the experimental data, also indicating that the available measurements are not suited to characterize the compressible behavior. In another study, the volumetric part of a two-term compressible Ogden model was included in the fitting of tension, compression and shear data of human brains and a good fit (*R*^2^ > 0.92) was achieved with an initial Poisson’s ratio of ≈ 0.42 (Moran et al., 2014). While these results first seem to be contradictory to others also achieving low fitting errors using quasi-incompressible formulations (MacManus et al., 2018; Pierrat et al., 2018; Shafieian et al., 2009; Hosseini-Farid et al., 2019), they are again explained by the lack of volumetric measurements that will cause non-unique solution if the compressible part is also fitted. Previous studies using image-based techniques come to the conclusion that the assumption of incompressible behavior can be justified for brain tissue (Felfelian et al., 2019; Eskandari et al., 2021).

Due to the aforementioned issues, we decided to predefine two different Poisson’s ratios and our results in Figure 13 show that a lower compressibility for a Poisson’s ratio of 0.49 comes along with significantly lower shear moduli than for 0.45, as the model yields higher forces for the same shear modulus and a higher bulk modulus under deformations causing volume changes, such as compression loading. As the obtained *α* values are mostly negative to capture the stiffer behavior in compression than in tension, the positive difference of the obtained values for a Poisson’s ratio of 0.49 in comparison to 0.45 mean that also the compression-stiffening behavior is less pronounced for a lower compressibility. Importantly, both parameter sets obtained by us using the Poisson’s ratios 0.45 and 0.49 are considered valid as they are able to replicate the measured experimental data. We therefore suggest to use the parameter set for a Poisson’s ratio of 0.49 representing quasi-incompressible behavior or, if the choice of the Poisson’s ratio is already fixed due to other constraints, the one with the closest value.

### Unconditioned and preconditioned material behavior

While we have only focused on the hyperelastic response of brain tissue in this study and have neglected poro- and viscoelastic effects, we have identified different parameter sets using the first and third cycle to represent the un- and preconditioned response, respectively. The results in Figure 14 and Supplementary Figure S11 show substantially lower shear moduli for experimental data from preconditioned specimens. This agrees well with previous studies highlighting the significant preconditioning effect of brain tissue (Budday et al., 2017a; Prevost et al., 2011; Labus et al., 2016). Although our statistical tests also confirm a significant difference in the nonlinearity parameter *α*, the low absolute values of the differences indicate a similar behavior in terms of the nonlinearity with the preconditioned parameters yielding only slightly more nonlinear behavior than the unconditioned parameters.

The appropriate parameter set for individual use cases can then be chosen based on the application. We note here, that the loading velocity of 40 μm/s (translating to a strain rate of ≈10^-2^ 1/s) in the experiments is comparably low. The obtained hyperelastic parameters are therefore approximating the equilibrium response. Thus, they are especially suited to model processes on the medium to longer time scale, such as brain growth and development or neurosurgery. The behavior of brain tissue in the aforementioned scenarios is best captured by the parameters using the preconditioned response. Nevertheless, if one wants to model impact scenarios related to, e.g., traumatic brain injury (TBI), and use a hyperelastic constitutive model, the parameters obtained from unconditioned data should be used. As the statistical analysis justifies the use of a constant *α* value, we provide the averaged values for the parameter sets obtained by fitting the averaged experimental response in different regions in Table 12.

**Table 12.**
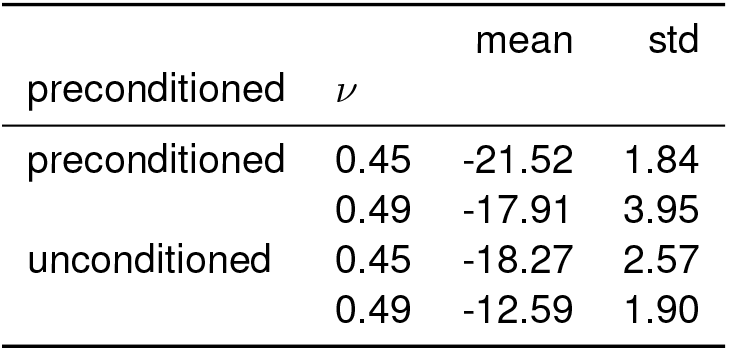
Averaged *α* parameter for the parameter sets fitted to the averaged experimental response in governing regions.

### Limitations

Its complex mechanical behavior and ultrasoft nature make the mechanical characterization of human brain tissue a challenging task. By choosing a hyperelastic model, we neglect the complex time-dependent behavior that can be observed in experiments (Budday et al., 2017b). An alternative are poro-viscoelastic models that capture also the biphasic nature of brain tissue (Greiner et al., 2021). However, currently the experimental data to accurately calibrate such complex models are not available and the assessment of regional trends – the main focus of this work – would be much more difficult. One of the pressing questions remaining for the mechanical characterization of brain tissue is to what extent parameters obtained from *ex vivo* tests can be applied to the prediction of the *in vivo* situation. In the latter, blood circulation and the restriction in its mobility by the bony skull capsule might play an important role. The differences in obtained material parameters between tested brains in Figure 11 and Supplementary Figure S7 were also confirmed as significant for single brains by the statistical tests. A potential reason for changes in properties could be metastases, as the cause of death for the concerned brains was metastasizing cancer. Still, other brains with the same cause of death did not show significantly different properties. Another reason could be the influence of different storage solutions as we obtained different nonlinearity parameters for the brain kept in Ringer’s solution. With the testing of more human brains in the future, it will be possible to better understand such effects, including person-specific data, such as age, sex, and cause of death.

A concern regarding parameter identification studies is the non-uniqueness of obtained solutions. Although we ran a global optimality study, shown in Supplementary Figure S14 to check if this behavior is observed in our case, this is just an empirical approach and not a rigorous mathematical analysis of the problem. Therefore, there is still the potential of occurring local minima for different experimental values.

Finally, to conclusively characterize the compressibility of human brain tissue, it would be necessary to capture the deformation of the specimens during testing, e.g. by using a camera. However, with the current testing setup, this is not easy to implement due to the bath of PBS required to control the temperature and hydration of the sample.

## Conclusion

The main goals of this work were to identify mechanically distinct regions in the human brain tissue and to subsequently provide region-specific parameters for finite element simulations. We tested human brain specimens under compression, tension and torsional shear in the finite strain regime. By fitting the response of a finite element model implementation of a modified one-term Ogden model to the experimental data, we have inversely identified material parameter sets. We have assigned the 19 anatomical regions from which the specimens were extracted to nine governing regions based on comparable microstructures and parameters. Statistical analyses show that at least the corona radiata and the corpus callosum should be modeled as mechanically distinct regions with different shear moduli. Interestingly the nonlinearity parameter *α* did not show significant differences. By fitting the first and third cycle of all loading modes separately, we have provided different parameter sets for the un- and preconditioned material responses, respectively. Varying the fixed initial Poisson’s ratio from 0.45 to 0.49 leads to a significant difference in the identified material parameters. In this respect, additional measurements are indespensible to reliably characterize the volumetric response of brain tissue in the future. In total, we have provided four parameter sets with distinct material parameters for the defined nine governing regions in Table 11 that can be used for mechanical simulations of the human brain. The most relevant parameter set can be selected based on the application at hand. Due to the relatively low loading velocity during our experiments, the parameter sets will be especially suited for mechanical models capturing effects that are occurring on a medium to long time scale.

## Acknowledgements

We would like to thank everyone who helped with the experiments and providing human brains, in particular, Lena Kretzschmar, Sarah Nistler, Jasmin Würges, Nicole Häck, and Lisa Stache. Moreover, the authors sincerely thank those who donated their bodies to science. Results from such research can potentially increase mankind’s overall knowledge that can then improve patient care. Therefore, these donors and their families deserve our highest gratitude.

## Declarations

### Ethical Approval

This study was approved by the Ethics Committee of Friedrich-Alexander-Universität Erlangen-Nürnberg, Germany, with the approval number 405_18 B. Body donors had given their written consent to donate their body to research.

### Authors’ contributions

S.B. conceptualized the study, acquired funding, and supervised the study. J.H. and S.B. developed the methodology. L.B. and F.P. provided human brains and expertise in human anatomy. N.R. performed the experiments and prepared the corresponding data. J.H. and S.K. implemented the computer code. J.H. performed all simulations, analyzed the data, and visualized the results. J.H., N.R., and S.B. wrote the initial manuscript draft. All authors discussed the results, reviewed and edited the manuscript.

### Availability of data and materials

The mechanical data and code generated and analyzed during the current study are available from the corresponding author on reasonable request.

## References

Arndt, Daniel, Wolfgang Bangerth, Denis Davydov, et al. (Jan. 2021). “The deal.II finite element library: Design, features, and insights”. In: Computers & Mathematics with Applications 81, pp. 407–422. DOI:10.1016/j.camwa.2020.02.022.

Blumcke, Ingmar, Silvia Budday, Annapurna Poduri, et al. (2021). “Neocortical development and epilepsy: insights from focal cortical dysplasia and brain tumours”. In: The Lancet Neurology 20.11, pp. 943–955.

Branch, Mary Ann, Thomas F. Coleman, and Yuying Li (Jan. 1999). “A Subspace, Interior, and >Conjugate Gradient Method for Large-Scale Bound-Constrained Minimization Problems”. In: SIAM Journal on Scientific Computing 21.1, pp. 1–23. DOI:10.1137/S1064827595289108.

Budday, S., G. Sommer, C. Birkl, et al. (Jan. 2017a). “Mechanical characterization of human brain tissue”. In: Acta Biomaterialia 48, pp. 319–340. DOI:10.1016/j.actbio.2016.10.036.

Budday, S., G. Sommer, J. Haybaeck, et al. (Sept. 2017b). “Rheological characterization of human brain tissue”. In: Acta Biomaterialia 60, pp. 315–329. DOI:10.1016/j.actbio.2017.06.024.

Budday, Silvia, Timothy C. Ovaert, Gerhard A. Holzapfel, et al. (July 2019). “Fifty Shades of Brain: A Review on the Mechanical Testing and Modeling of Brain Tissue”. In: Archives of Computational Methods in Engineering. DOI:10.1007/s11831-019-09352-w.

Budday, Silvia, Paul Steinmann, and Ellen Kuhl (July 2015). “Physical biology of human brain development”. In: Frontiers in Cellular Neuroscience 9. DOI:10.3389/fncel.2015.00257.

Chatelin, S., J. Vappou, S. Roth, et al. (Feb. 2012). “Towards child versus adult brain mechanical properties”. In: Journal of the Mechanical Behavior of Biomedical Materials 6, pp. 166–173. DOI:10.1016/j.jmbbm.2011.09.013.

Delorme, Sébastien, Denis Laroche, Robert DiRaddo, et al. (Sept. 2012). “NeuroTouch”. In: Operative Neurosurgery 71.suppl_1, ons32–ons42. DOI:10.1227/NEU.0b013e318249c744.

Eskandari, Faezeh, Zahra Rahmani, and Mehdi Shafieian (2021). “The effect of large deformation on Poisson’s ratio of brain white matter: An experimental study”. In: Proceedings of the Institution of Mechanical Engineers, Part H: Journal of Engineering in Medicine 235.4, pp. 401–407. DOI:10.1177/0954411920984027.

Faber, Jessica, Jan Hinrichsen, Alexander Greiner, et al. (Apr. 2022). “Tissue-Scale Biomechanical Testing of Brain Tissue for the Calibration of Nonlinear Material Models”. In: Current Protocols 2.4. DOI:10.1002/cpz1.381.

Felfelian, Amir Mohammad, Amirhosein Baradaran Najar, Reza Jafari Nedoushan, et al. (Dec. 2019). “Determining constitutive behavior of the brain tissue using digital image correlation and finite element modeling”. In: Biomechanics and Modeling in Mechanobiology 18.6, pp. 1927–1945. DOI:10.1007/s10237-019-01186-6.

Feng, Yuan, Chung-Hao Lee, Lining Sun, et al. (Jan. 2017). “Characterizing white matter tissue in large strain via asymmetric indentation and inverse finite element modeling”. In: Journal of the Mechanical Behavior of Biomedical Materials 65, pp. 490–501. DOI: 10.1016/j.jmbbm.2016.09.020.

Fernandes, Fábio A.O., Dmitri Tchepel, Ricardo J. Alves de Sousa, et al. (Jan. 2018). “Development and validation of a new finite element human head model: Yet another head model (YEAHM)”. In: Engineering Computations 35.1, pp. 477–496. DOI:10.1108/EC-09-2016-0321.

Finan, John D., Sowmya N. Sundaresh, Benjamin S. Elkin, et al. (June 2017). “Regional mechanical properties of human brain tissue for computational models of traumatic brain injury”. In: Acta Biomaterialia 55, pp. 333–339. DOI:10.1016/j.actbio.2017.03.037.

Forte, Antonio E., Stephen M. Gentleman, and Daniele Dini (June 2017). “On the characterization of the heterogeneous mechanical response of human brain tissue”. In: Biomechanics and Modeling in Mechanobiology 16.3, pp. 907–920. DOI:10.1007/s10237-016-0860-8.

Garcia, KE, CD Kroenke, and PV Bayly (2018). “Mechanics of cortical folding: stress, growth and stability”. In: Philosophical Transactions of the Royal Society B: Biological Sciences 373.1759, p. 20170321.

Gavrus, A., E. Massoni, and J.L. Chenot (June 1996). “An inverse analysis using a finite element model for identification of rheological parameters”. In: Journal of Materials Processing Technology 60.1-4, pp. 447–454. DOI:10.1016/0924-0136(96)02369-2.

Ghajari, Mazdak, Peter J. Hellyer, and David J. Sharp (Feb. 2017). “Computational modelling of traumatic brain injury predicts the location of chronic traumatic encephalopathy pathology”. In: Brain 140.2, pp. 333–343. DOI:10.1093/brain/aww317.

Giudice, J. Sebastian, Ahmed Alshareef, Taotao Wu, et al. (2021). “Calibration of a Heterogeneous Brain Model Using a Subject-Specific Inverse Finite Element Approach”. In: Frontiers in Bioengineering and Biotechnology 9.

Greiner, Alexander, Nina Reiter, Friedrich Paulsen, et al. (Aug. 2021). “Poro-Viscoelastic Effects During Biomechanical Testing of Human Brain Tissue”. In: Frontiers in Mechanical Engineering 7, p. 708350. DOI:10.3389/fmech.2021.708350.

Hansen, Kim V., Lars Brix, Christian F. Pedersen, et al. (Mar. 2004). “Modelling of interaction between a spatula and a human brain”. In: Medical Image Analysis 8.1, pp. 23–33. DOI: 10.1016/j.media.2003.07.001.

Ho, Johnson and Svein Kleiven (2009). “Can sulci protect the brain from traumatic injury?” In: Journal of Biomechanics 42.13, pp. 2074–2080. DOI:https://doi.org/10.1016/j.jbiomech.2009.06.051.

Holzapfel, Gerhard A. (2000). Nonlinear solid mechanics: a continuum approach for engineering. Chichester; New York: Wiley.

Horgan, T J and M D Gilchrist (Jan. 2003). “The creation of three-dimensional finite element models for simulating head impact biomechanics”. In: International Journal of Crashworthiness 8.4, pp. 353–366. DOI:10.1533/ijcr.2003.0243.

Hosseini-Farid, Mohammad, Mohammadreza Ramzanpour, Mariusz Ziejewski, et al. (Nov. 2019). “A compressible hyper-viscoelastic material constitutive model for human brain tissue and the identification of its parameters”. In: International Journal of Non-Linear Mechanics 116, pp. 147–154. DOI:10.1016/j.ijnonlinmec.2019.06.008.

Hunter, John D. (May 2007). “Matplotlib: A 2D Graphics Environment”. In: Computing in Science & Engineering 9.3, pp. 90–95. DOI:10.1109/MCSE.2007.55.

Ji, Songbai, Mazdak Ghajari, Haojie Mao, et al. (July 2022). “Use of Brain Biomechanical Models for Monitoring Impact Exposure in Contact Sports”. In: Annals of Biomedical Engineering. DOI:10.1007/s10439-022-02999-w.

Jin, Xin, Feng Zhu, Haojie Mao, et al. (Nov. 2013). “A comprehensive experimental study on material properties of human brain tissue”. In: Journal of Biomechanics 46.16, pp. 2795–2801. DOI:10.1016/j.jbiomech.2013.09.001.

Kang, Ho-Sung, Rémy Willinger, Baye M. Diaw, et al. (Nov. 1997). Validation of a 3D Anatomic Human Head Model and Replication of Head Impact in Motorcycle Accident by Finite Element Modeling. SAE Technical Paper 973339. Warrendale, PA: SAE International. DOI: 10.4271/973339.

Karimi, Alireza, Seyed Mohammadali Rahmati, Reza Razaghi, et al. (Jan. 2019). “Mechanical measurement of the human cerebellum under compressive loading”. In: Journal of Medical Engineering & Technology 43.1, pp. 55–58. DOI:10.1080/03091902.2019.1609609.

King, Andrew and Robert Eckersley (2019). Statistics for Biomedical Engineers and Scientists. Elsevier. DOI:10.1016/C2018-0-02241-0.

Labus, Kevin M. and Christian M. Puttlitz (Sept. 2016). “An anisotropic hyperelastic constitutive model of brain white matter in biaxial tension and structural–mechanical relationships”. In: Journal of the Mechanical Behavior of Biomedical Materials 62, pp. 195–208. DOI:10.1016/j.jmbbm.2016.05.003.

Laksari, Kaveh, Mehdi Shafieian, and Kurosh Darvish (Feb. 2012). “Constitutive model for brain tissue under finite compression”. In: Journal of Biomechanics 45.4, pp. 642–646. DOI:10.1016/j.jbiomech.2011.12.023.

Li, Xiaogai, Zhou Zhou, and Svein Kleiven (2021). “An anatomically detailed and personalizable head injury model: Significance of brain and white matter tract morphological variability on strain”. In: Biomechanics and modeling in mechanobiology 20.2, pp. 403–431.

MacManus, David B., Andrea Menichetti, Bart Depreitere, et al. (Nov. 2020). “Towards animal surrogates for characterising large strain dynamic mechanical properties of human brain tissue”. In: Brain Multiphysics 1, p. 100018. DOI:10.1016/j.brain.2020.100018.

MacManus, David B., Jeremiah G. Murphy, and Michael D. Gilchrist (Nov. 2018). “Mechanical characterisation of brain tissue up to 35% strain at 1, 10, and 100/s using a custom-built micro-indentation apparatus”. In: Journal of the Mechanical Behavior of Biomedical Materials 87, pp. 256–266. DOI:10.1016/j.jmbbm.2018.07.025.

Majdan, Marek, Dominika Plancikova, Andrew Maas, et al. (July 2017). “Years of life lost due to traumatic brain injury in Europe: A cross-sectional analysis of 16 countries”. In: PLOS Medicine 14.7. Ed. by Martin Schreiber, e1002331. DOI:10.1371/journal.pmed.1002331.

Mao, Haojie, Liying Zhang, Binhui Jiang, et al. (Sept. 2013). “Development of a Finite Element Human Head Model Partially Validated With Thirty Five Experimental Cases”. In: Journal of Biomechanical Engineering 135.11. DOI:10.1115/1.4025101.

McKinney, Wes (2010). “Data Structures for Statistical Computing in Python”. In: vol. 445. Austin, Texas, pp. 56–61. DOI:10.25080/Majora-92bf1922-00a.

Menichetti, Andrea, David B. MacManus, Michael D. Gilchrist, et al. (Oct. 2020). “Regional characterization of the dynamic mechanical properties of human brain tissue by microindentation”. In: International Journal of Engineering Science 155, p. 103355. DOI:10.1016/j.ijengsci.2020.103355.

Mihai, L. Angela, Silvia Budday, Gerhard A. Holzapfel, et al. (Sept. 2017a). “A family of hyperelastic models for human brain tissue”. In: Journal of the Mechanics and Physics of Solids 106, pp. 60–79. DOI:10.1016/j.jmps.2017.05.015.

Mihai, L. Angela, LiKang Chin, Paul A. Janmey, et al. (Sept. 2015). “A comparison of hyperelastic constitutive models applicable to brain and fat tissues”. In: Journal of The Royal Society Interface 12.110, p. 20150486. DOI:10.1098/rsif.2015.0486.

Mihai, L. Angela and Alain Goriely (Nov. 2017b). “How to characterize a nonlinear elastic material? A review on nonlinear constitutive parameters in isotropic finite elasticity”. In: Proceedings of the Royal Society A: Mathematical, Physical and Engineering Sciences 473.2207, p. 20170607. DOI:10.1098/rspa.2017.0607.

Miller, Karol (Jan. 2005). “Method of testing very soft biological tissues in compression”. In: Journal of Biomechanics 38.1, pp. 153–158. DOI:10.1016/j.jbiomech.2004.03.004.

Miller, Karol, Kiyoyuki Chinzei, Girma Orssengo, et al. (Nov. 2000). “Mechanical properties of brain tissue in-vivo: experiment and computer simulation”. In: Journal of Biomechanics 33.11, pp. 1369–1376. DOI:10.1016/S0021-9290(00)00120-2.

Moran, Richard, Joshua H. Smith, and José J. García (Nov. 2014). “Fitted hyperelastic parameters for Human brain tissue from reported tension, compression, and shear tests”. In: Journal of Biomechanics 47.15, pp. 3762–3766. DOI:10.1016/j.jbiomech.2014.09.030.

Nair, Sudhakar (2009). Introduction to continuum mechanics. New York, NY: Cambridge University Press.

Nelder, J. A. and R. Mead (Jan. 1965). “A Simplex Method for Function Minimization”. In: The Computer Journal 7.4, pp. 308–313. DOI:10.1093/comjnl/7.4.308.

Ogden, R. W. (June 1972). “Large deformation isotropic elasticity: on the correlation of theory and experiment for compressible rubberlike solids”. In: Proceedings of the Royal Society of London. A. Mathematical and Physical Sciences 328.1575, pp. 567–583. DOI:10.1098/rspa.1972.0096.

Pierrat, B., D.B. MacManus, J.G. Murphy, et al. (Feb. 2018). “Indentation of heterogeneous soft tissue: Local constitutive parameter mapping using an inverse method and an automated rig”. In: Journal of the Mechanical Behavior of Biomedical Materials 78, pp. 515–528. DOI:10.1016/j.jmbbm.2017.03.033.

Prevost, Thibault P., Asha Balakrishnan, Subra Suresh, et al. (Jan. 2011). “Biomechanics of brain tissue”. In: Acta Biomaterialia 7.1, pp. 83–95. DOI:10.1016/j.actbio.2010.06.035.

Sase, Kazuya, Akira Fukuhara, Teppei Tsujita, et al. (Dec. 2015). “GPU-accelerated surgery simulation for opening a brain fissure”. In: ROBOMECH Journal 2.1. DOI:10.1186/s40648-015-0040-0.

Seber, George A. F and C. J Wild (2005). Nonlinear Regression. Wiley series in probability and mathematical statistics. Probability and mathematical statistics. Hoboken, NJ: Wiley.

Shafieian, Mehdi, Kurosh K. Darvish, and James R. Stone (Sept. 2009). “Changes to the viscoelastic properties of brain tissue after traumatic axonal injury”. In: Journal of Biomechanics 42.13, pp. 2136–2142. DOI:10.1016/j.jbiomech.2009.05.041.

Simo, J.C. and C. Miehe (July 1992). “Associative coupled thermoplasticity at finite strains: Formulation, numerical analysis and implementation”. In: Computer Methods in Applied Mechanics and Engineering 98.1, pp. 41–104.DOI:10.1016/0045-7825(92)90170-0.

Terpilowski, Maksim (Apr. 2019). “scikit-posthocs: Pairwise multiple comparison tests in Python”. In: Journal of Open Source Software 4.36, p. 1169. DOI:10.21105/joss.01169.

Virtanen, Pauli, Ralf Gommers, Travis E. Oliphant, et al. (Mar. 2020). “SciPy 1.0: fundamental algorithms for scientific computing in Python”. In: Nature Methods 17.3, pp. 261–272. DOI:10.1038/s41592-019-0686-2.

Voyiadjis, George Z. and Aref Samadi-Dooki (July 2018). “Hyperelastic modeling of the human brain tissue: Effects of no-slip boundary condition and compressibility on the uniaxial deformation”. In: Journal of the Mechanical Behavior of Biomedical Materials 83, pp. 63–78. DOI:10.1016/j.jmbbm.2018.04.011.

Waskom, Michael (Apr. 2021). “seaborn: statistical data visualization”. In: Journal of Open Source Software 6.60, p. 3021. DOI:10.21105/joss.03021.

Weickenmeier, Johannes, Ellen Kuhl, and Alain Goriely (2018). “Multiphysics of prionlike diseases: Progression and atrophy”. In: Physical review letters 121.15, p. 158101.

Wu, Taotao, Ahmed Alshareef, J. Sebastian Giudice, et al. (Sept. 2019). “Explicit Modeling of White Matter Axonal Fiber Tracts in a Finite Element Brain Model”. In: Annals of Biomedical Engineering 47.9, pp. 1908–1922. DOI:10.1007/s10439-019-02239-8.

Yan, Wenyi and Oscar Dwiputra Pangestu (Dec. 2011). “A modified human head model for the study of impact head injury”. In: Computer Methods in Biomechanics and Biomedical Engineering 14.12, pp. 1049–1057. DOI:10.1080/10255842.2010.506435.

Zarzor, M.S., S. Kaessmair, P. Steinmann, et al. (2021). “A two-field computational model couples cellular brain development with cortical folding”. In: Brain Multiphysics 2, p. 100025. DOI:10.1016/j.brain.2021.100025.

Zhao, Wei and Songbai Ji (Feb. 2022). “Cerebral vascular strains in dynamic head impact using an upgraded model with brain material property heterogeneity”. In: Journal of the Mechanical Behavior of Biomedical Materials 126, p. 104967. DOI:10.1016/j.jmbbm.2021.104967.

Zhu, Feng, Xin Jin, Fengjiao Guan, et al. (Dec. 2010). “Identifying the properties of ultra-soft materials using a new methodology of combined specimen-specific finite element model and optimization techniques”. In: Materials & Design 31.10, pp. 4704–4712. DOI:10.1016/j.matdes.2010.05.023.

Zong, Z., H. P. Lee, and C. Lu (Jan. 2006). “A three-dimensional human head finite element model and power flow in a human head subject to impact loading”. In: Journal of Biomechanics 39.2, pp. 284–292. DOI:10.1016/j.jbiomech.2004.11.015.

