## Supplementary Material for "Inverse identification of region-specific hyperelastic material parameters for human brain tissue"

#### 1 Parameter sets

| | preconditioned $\nu$ =45 | | | | preconditioned $\nu$ =49 | | | | unconditioned $\nu$ =45 | | | | unconditioned $\nu$ =49 | | | |
|---|---|---|---|---|---|---|---|---|---|---|---|---|---|---|---|---|
| | $\alpha$ | | $\mu$ | | $\alpha$ | | $\mu$ | | $\alpha$ | | $\mu$ | | $\alpha$ | | $\mu$ | |
| | mean | std. | mean | std. | mean | std. | mean | std. | mean | std. | mean | std. | mean | std. | mean | std. |
| gov. region | | | | | | | | | | | | | | | | |
| Am | -19.67 | 1.80 | 257.05 | 116.18 | -14.16 | 1.60 | 193.34 | 100.33 | -14.74 | 2.55 | 386.97 | 171.00 | -10.10 | 2.26 | 293.06 | 133.23 |
| BG | -18.95 | 4.02 | 252.15 | 99.15 | -15.05 | 4.80 | 169.67 | 66.85 | -15.90 | 4.51 | 338.67 | 135.90 | -11.49 | 4.14 | 246.21 | 97.77 |
| BS | -20.41 | 6.47 | 233.12 | 112.45 | -19.59 | 8.21 | 118.06 | 43.22 | -16.82 | 6.11 | 336.44 | 165.75 | -13.60 | 5.99 | 225.08 | 106.82 |
| C | -19.27 | 4.42 | 281.75 | 114.14 | -16.72 | 5.05 | 175.21 | 73.39 | -16.16 | 2.92 | 392.24 | 153.61 | -12.16 | 2.79 | 285.97 | 110.82 |
| CB | -23.22 | 5.96 | 177.49 | 81.38 | -18.79 | 3.76 | 115.36 | 48.54 | -19.49 | 4.03 | 255.38 | 100.99 | -14.08 | 3.40 | 196.55 | 88.38 |
| CC | -14.94 | 12.60 | 61.66 | 24.73 | -13.72 | 13.71 | 37.64 | 18.26 | -12.82 | 7.88 | 95.43 | 34.55 | -9.11 | 6.90 | 67.82 | 24.40 |
| CR | -19.70 | 4.73 | 157.09 | 75.55 | -19.08 | 7.60 | 87.58 | 47.53 | -16.93 | 6.60 | 218.75 | 115.76 | -13.55 | 6.94 | 146.54 | 77.74 |
| Hi | -17.37 | 2.75 | 184.21 | 81.38 | -13.09 | 3.15 | 129.96 | 47.59 | -13.25 | 1.58 | 292.58 | 147.86 | -9.23 | 1.33 | 218.74 | 112.20 |
| M | -18.68 | 4.99 | 215.70 | 74.61 | -15.15 | 5.96 | 137.67 | 37.76 | -15.11 | 4.97 | 314.92 | 110.47 | -10.89 | 4.19 | 227.69 | 74.32 |

Table S1: Material parameters averaged over governing regions.

| | preconditioned $\nu$ =45 | | | | preconditioned $\nu$ =49 | | | | unconditioned $\nu$ =45 | | | | unconditioned $\nu$ =49 | | | |
|---|---|---|---|---|---|---|---|---|---|---|---|---|---|---|---|---|
| | $\alpha$ | | $\mu$ | | $\alpha$ | | $\mu$ | | $\alpha$ | | $\mu$ | | $\alpha$ | | $\mu$ | |
| | mean | std. | mean | std. | mean | std. | mean | std. | mean | std. | mean | std. | mean | std. | mean | std. |
| region | | | | | | | | | | | | | | | | |
| Am | -19.67 | 1.80 | 257.05 | 116.18 | -14.16 | 1.60 | 193.34 | 100.33 | -14.74 | 2.55 | 386.97 | 171.00 | -10.10 | 2.26 | 293.06 | 133.23 |
| CC | -14.94 | 12.60 | 61.66 | 24.73 | -13.72 | 13.71 | 37.64 | 18.26 | -12.82 | 7.88 | 95.43 | 34.55 | -9.11 | 6.90 | 67.82 | 24.40 |
| CI | -21.20 | - | 205.14 | - | -19.49 | - | 116.44 | - | -18.37 | - | 261.34 | - | -13.99 | - | 188.13 | - |
| CR | -19.83 | 2.96 | 155.69 | 67.08 | -19.28 | 5.99 | 83.33 | 27.13 | -18.65 | 2.02 | 209.72 | 97.64 | -15.78 | 3.89 | 136.21 | 57.59 |
| FC | -19.63 | 0.97 | 249.69 | 82.67 | -15.90 | 1.90 | 168.72 | 53.82 | -15.29 | 1.73 | 361.23 | 127.01 | -11.08 | 1.75 | 271.89 | 96.14 |
| Hi | -17.37 | 2.75 | 184.21 | 81.38 | -13.09 | 3.15 | 129.96 | 47.59 | -13.25 | 1.58 | 292.58 | 147.86 | -9.23 | 1.33 | 218.74 | 112.20 |
| M | -18.72 | 4.87 | 230.60 | 99.27 | -16.35 | 6.36 | 135.61 | 42.66 | -15.07 | 4.16 | 338.29 | 137.77 | -11.30 | 4.13 | 238.30 | 91.76 |
| MC | -20.19 | 1.64 | 318.98 | 126.18 | -17.99 | 3.06 | 195.50 | 84.99 | -16.95 | 2.06 | 434.35 | 168.54 | -12.94 | 2.19 | 313.65 | 119.22 |
| Me | -18.90 | 7.63 | 228.86 | 106.41 | -17.96 | 9.30 | 120.53 | 41.93 | -16.11 | 7.20 | 323.46 | 161.20 | -13.15 | 6.92 | 216.22 | 107.02 |
| NC | -15.82 | 5.85 | 198.57 | 99.88 | -11.39 | 5.73 | 143.46 | 74.60 | -12.58 | 6.10 | 277.58 | 138.29 | -8.30 | 4.94 | 203.23 | 96.48 |
| P | -23.23 | 1.31 | 241.04 | 131.47 | -22.62 | 4.91 | 113.46 | 48.61 | -18.12 | 3.38 | 360.56 | 184.33 | -14.43 | 4.09 | 241.52 | 112.86 |
| Pa | -21.16 | 2.14 | 242.06 | 76.14 | -17.94 | 3.75 | 154.22 | 60.56 | -18.19 | 3.07 | 332.32 | 137.93 | -13.67 | 3.21 | 240.96 | 105.92 |
| Pu | -19.24 | 1.76 | 305.10 | 100.76 | -15.09 | 2.90 | 206.07 | 55.33 | -16.27 | 2.61 | 393.89 | 121.48 | -11.86 | 2.75 | 285.82 | 82.46 |
| TL | -8.37 | 13.00 | 123.68 | 23.29 | -4.38 | 11.03 | 89.51 | 23.47 | -9.18 | 3.07 | 202.19 | 75.27 | -5.95 | 1.97 | 150.13 | 61.67 |
| Th | -18.66 | 5.27 | 205.38 | 53.76 | -14.32 | 5.78 | 139.10 | 35.73 | -15.13 | 5.63 | 298.74 | 89.50 | -10.60 | 4.38 | 220.34 | 62.52 |
| VC | -20.81 | 0.58 | 299.02 | 67.23 | -19.10 | 1.09 | 174.95 | 47.48 | -17.64 | 1.70 | 421.88 | 107.24 | -13.62 | 1.80 | 306.03 | 90.17 |
| WM | -19.62 | 5.66 | 158.00 | 82.06 | -18.95 | 8.62 | 90.34 | 57.54 | -15.81 | 8.21 | 224.64 | 127.98 | -12.10 | 8.11 | 153.27 | 89.07 |
| cN | -22.66 | 4.42 | 123.64 | 38.58 | -17.43 | 2.83 | 89.13 | 39.37 | -17.50 | 2.53 | 203.00 | 84.30 | -12.70 | 2.90 | 155.49 | 74.71 |
| cWM | -23.57 | 7.03 | 211.15 | 84.48 | -19.65 | 4.18 | 131.76 | 48.53 | -20.74 | 4.43 | 288.12 | 101.18 | -14.94 | 3.57 | 222.22 | 90.80 |

Table S2: Material parameters averaged over anatomical regions.

#### 2 Normality

| $\nu$ | precond | RMSE | | $\alpha$ | | $\mu$ | |
|---|---|---|---|---|---|---|---|
| | | W | p | W | p | W | p |
| 0.45 | preconditioned | 0.861346 | 7.426005e-12 | 7.35e-01 | 9.06e-17 | 9.61e-01 | 5.68e-05 |
| 0.45 | unconditioned | 0.843182 | 1.023565e-12 | 8.29e-01 | 2.53e-13 | 9.57e-01 | 2.38e-05 |
| 0.49 | preconditioned | 0.821075 | 1.123478e-13 | 9.29e-01 | 8.81e-08 | 9.46e-01 | 2.15e-06 |
| 0.49 | unconditioned | 0.844200 | 1.138903e-12 | 8.90e-01 | 2.43e-10 | 9.54e-01 | 1.24e-05 |

Table S3: Results for the Shapiro Wilk test for the material parameters $\mu$ and $\alpha$ yielding the probability that the observations stem from a normal distribution.

#### 3 Regional dependency

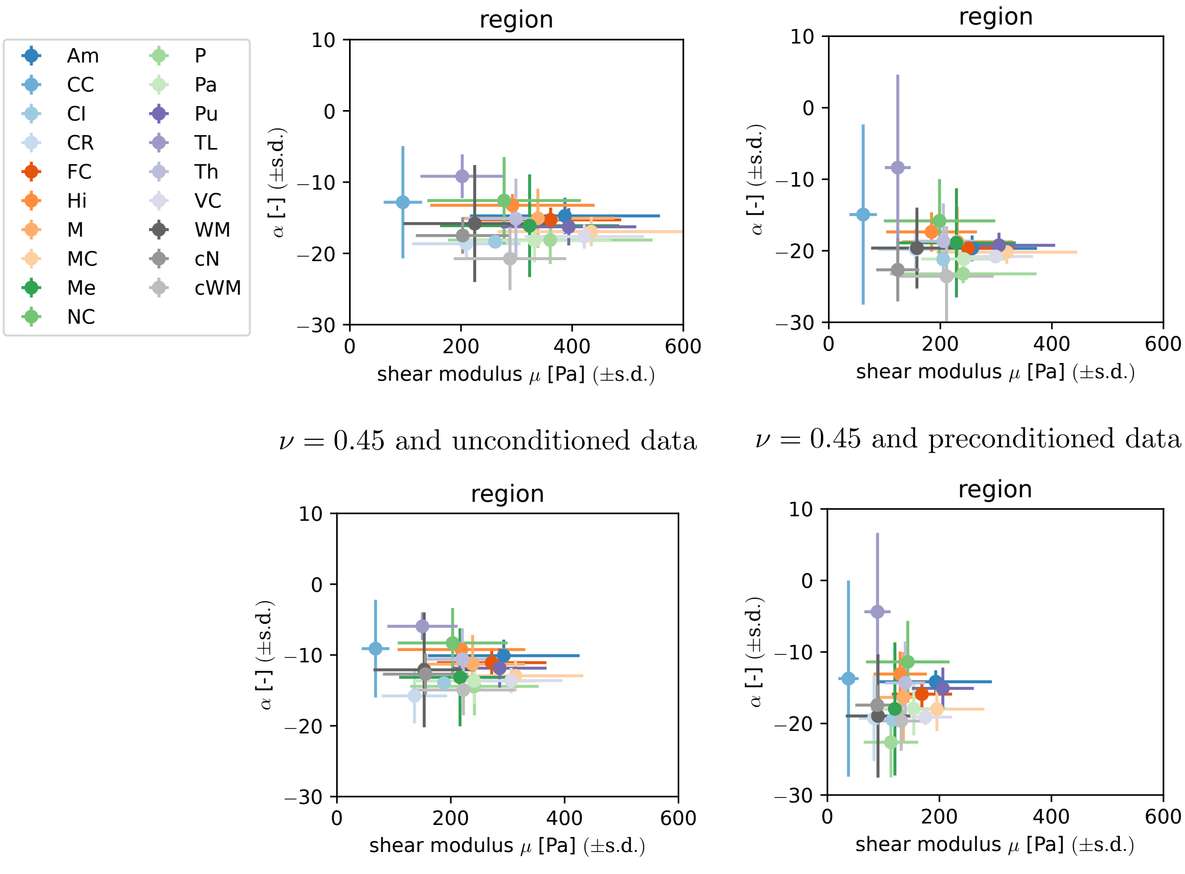


$\nu = 0.45$ and unconditioned data $\nu = 0.45$ and preconditioned data

$\nu = 0.49$ and unconditioned data $\nu = 0.49$ and preconditioned data

Figure S1: Material parameter distribution for different anatomical regions.

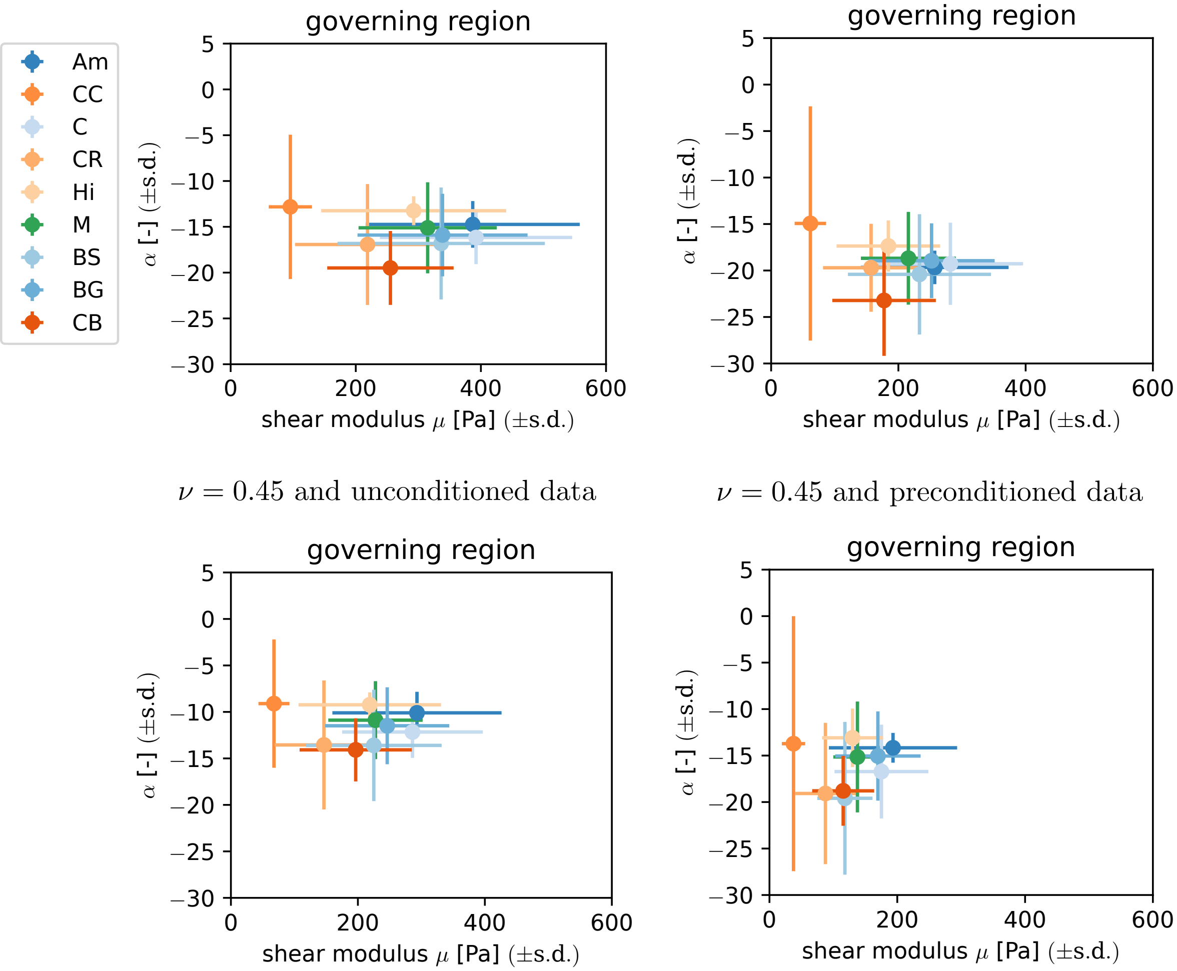


$\nu = 0.45$ and unconditioned data

$\nu = 0.45$ and preconditioned data

$\nu = 0.49$ and unconditioned data

$\nu = 0.49$ and preconditioned data

Figure S2: Material parameter distribution for the defined governing regions.

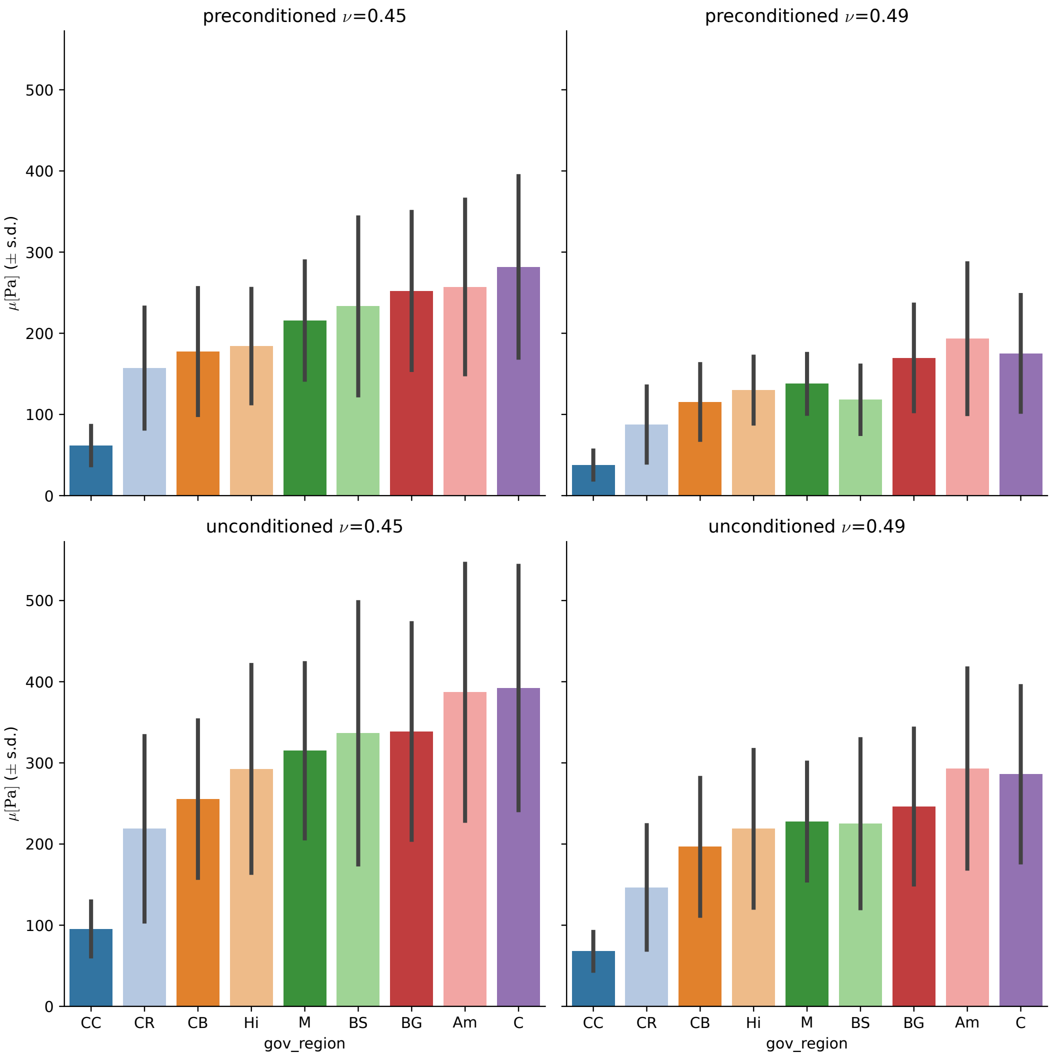


Figure S3: Resulting values for the shear modulus $\mu$, averaged over the governing regions.

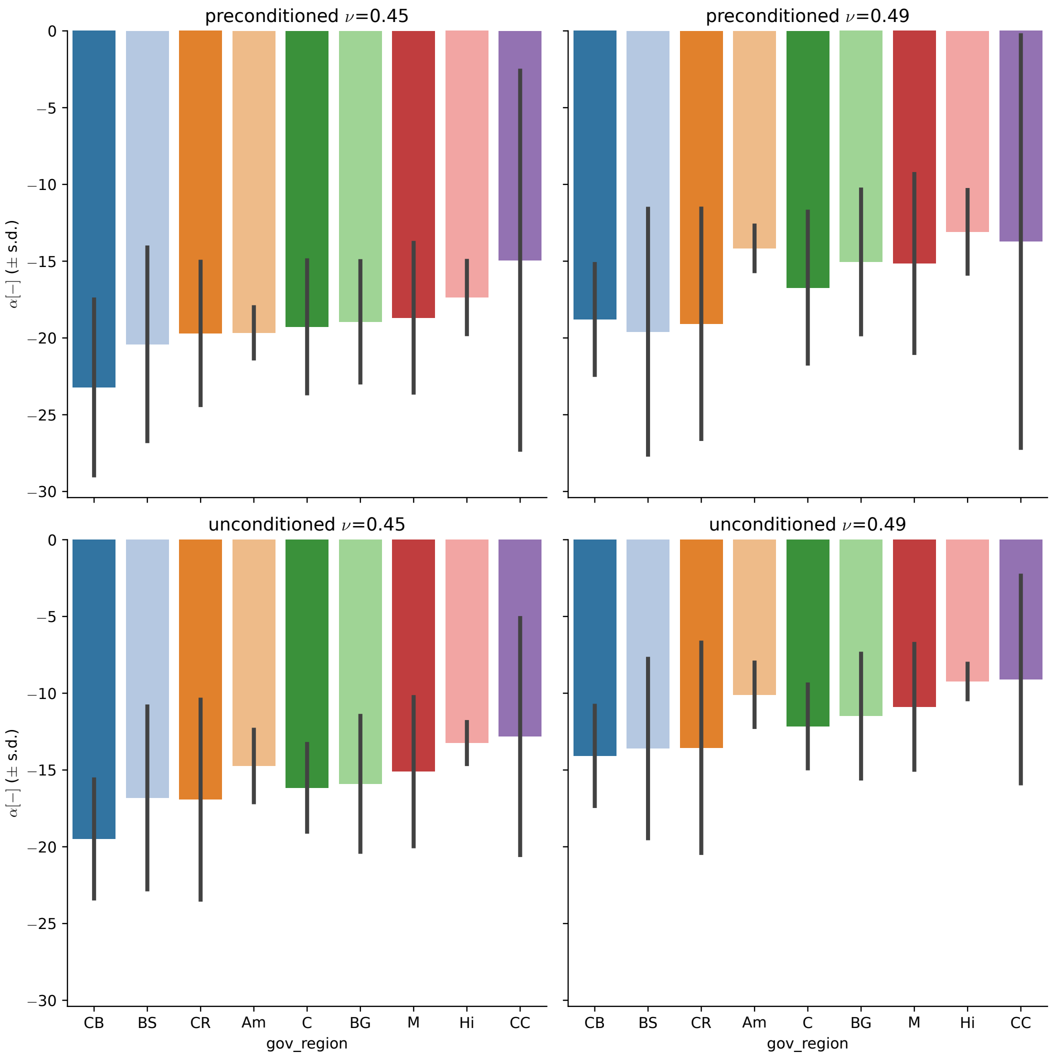


Figure S4: Resulting values for the nonlinearity parameter $\alpha$, averaged over the governing regions.

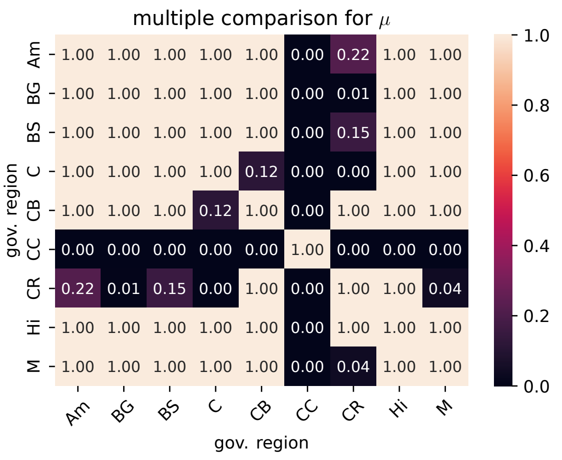


$\nu = 0.45$ and unconditioned data

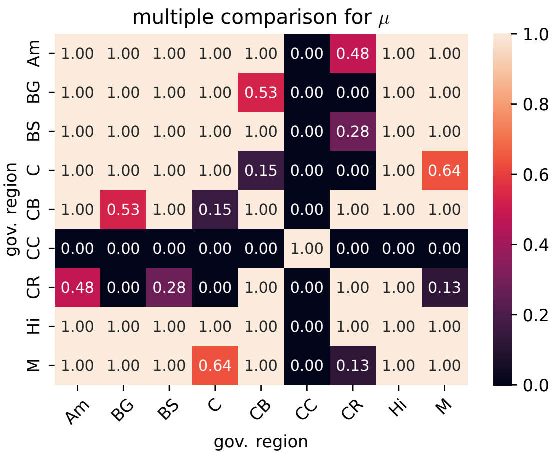


$\nu = 0.45$ and preconditioned data

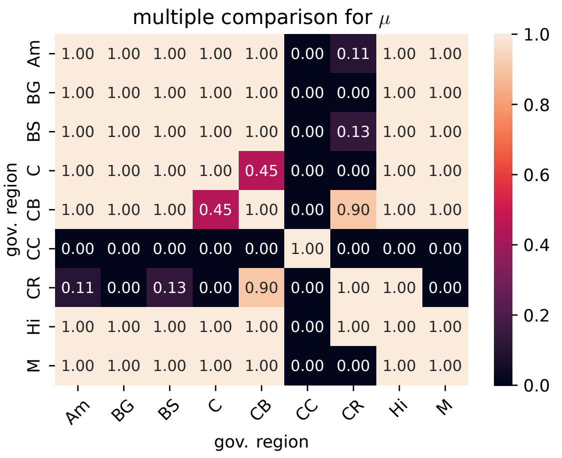


$\nu = 0.49$ and unconditioned data

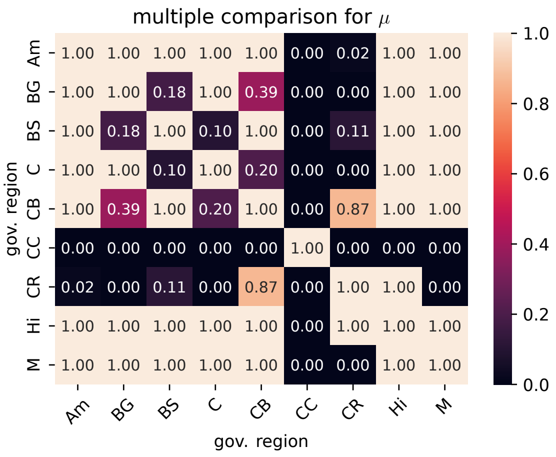


$\nu = 0.49$ and preconditioned data

Figure S5: Resulting $p$-values from pairwise post hoc Mann-Whitney-U tests comparing the shear modulus $\mu$ for the different governing regions. Low $p$ values indicate that the corpus callosum and the corona radiata have distinct properties.

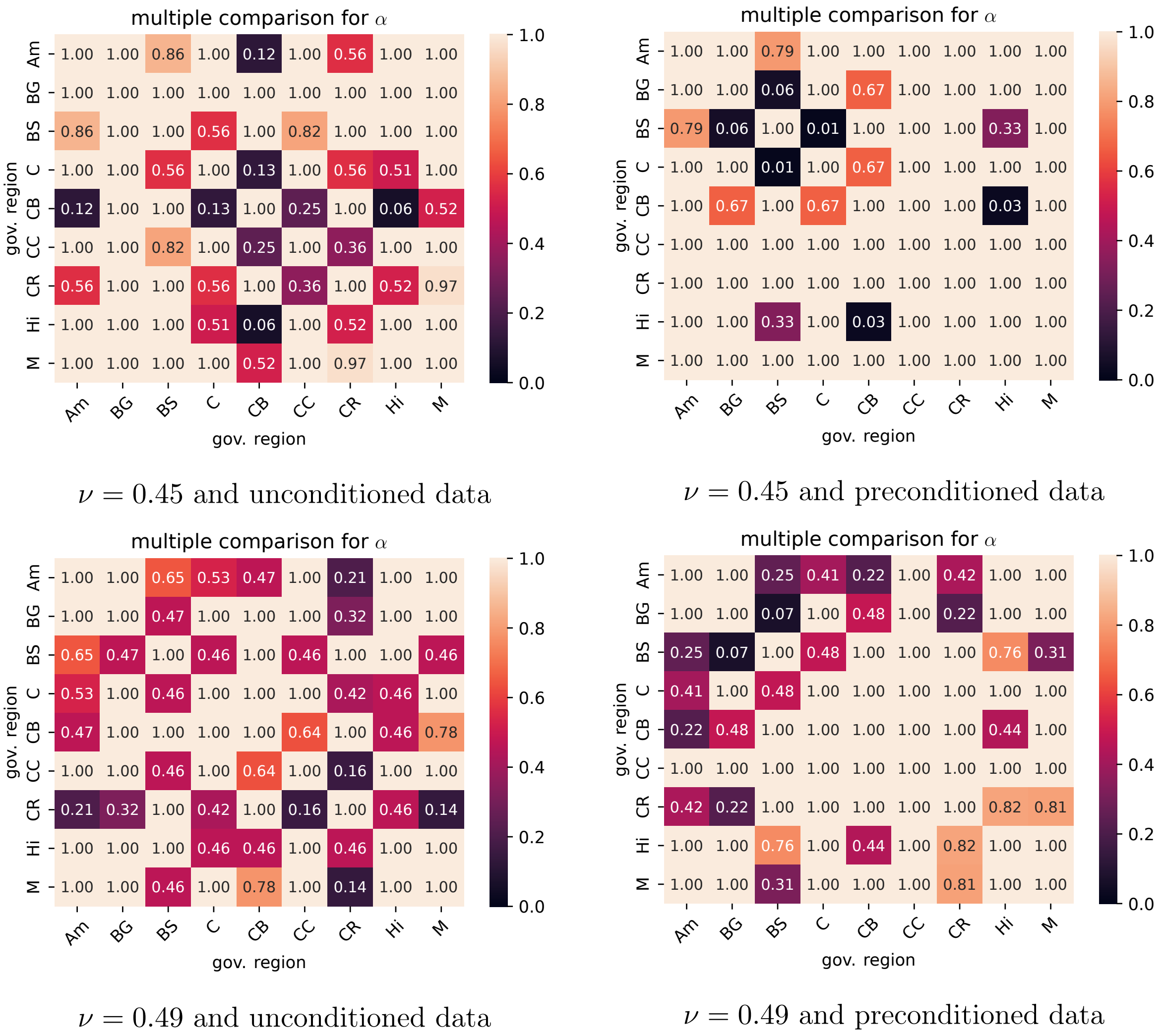


$\nu = 0.45$ and unconditioned data

$\nu = 0.45$ and preconditioned data

$\nu = 0.49$ and unconditioned data

$\nu = 0.49$ and preconditioned data

Figure S6: Resulting $p$-values from pairwise post hoc Mann-Whitney-U tests comparing the nonlinearity parameter $\alpha$ for the different governing regions. A lack of low $p$ values indicates that no distinct regions are found.

#### 4 Inter-individual variation

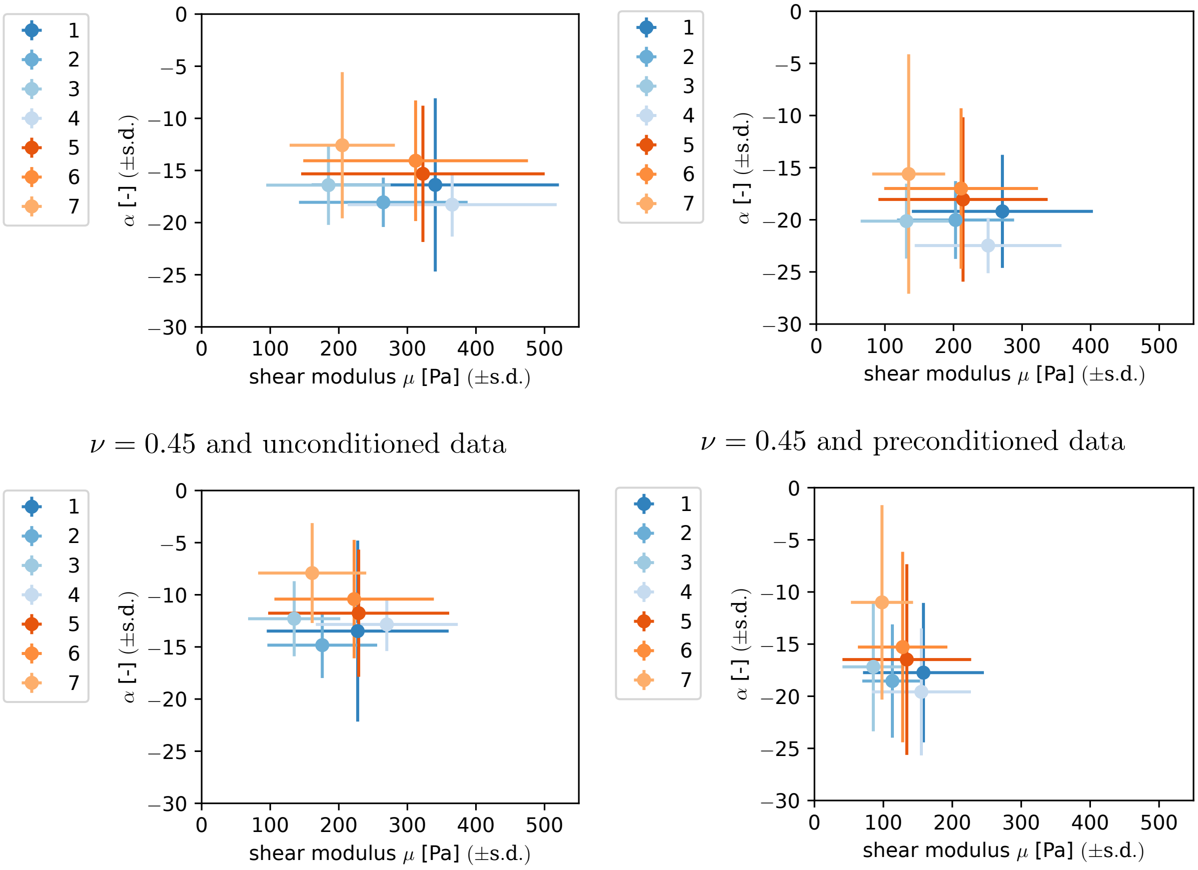


$\nu = 0.45$ and unconditioned data

$\nu = 0.45$ and preconditioned data

$\nu = 0.49$ and unconditioned data

$\nu = 0.49$ and preconditioned data

Figure S7: Parameter values and their standard deviation for different tested brains.

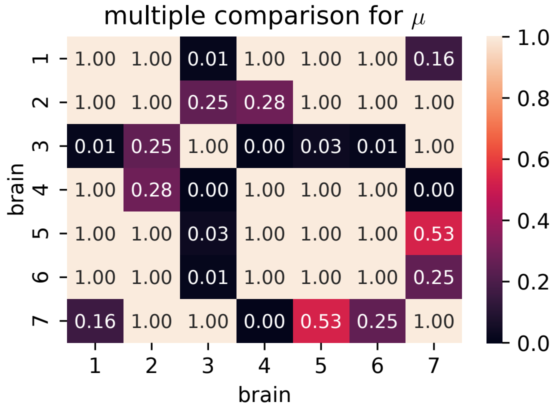


$\nu = 0.45$ and unconditioned data

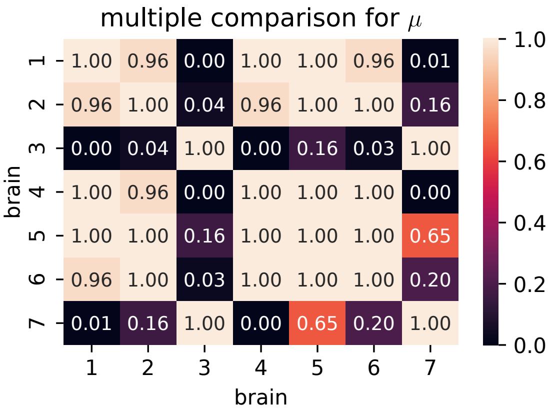


$\nu = 0.45$ and preconditioned data

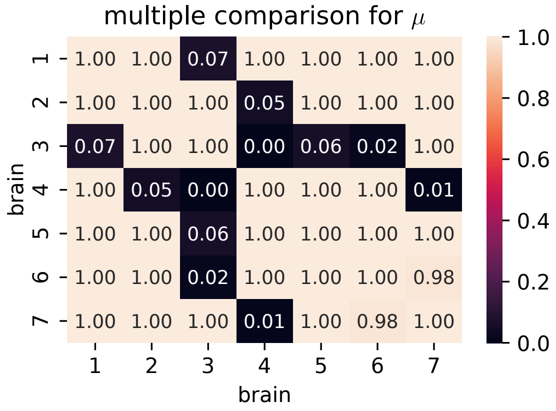


$\nu = 0.49$ and unconditioned data

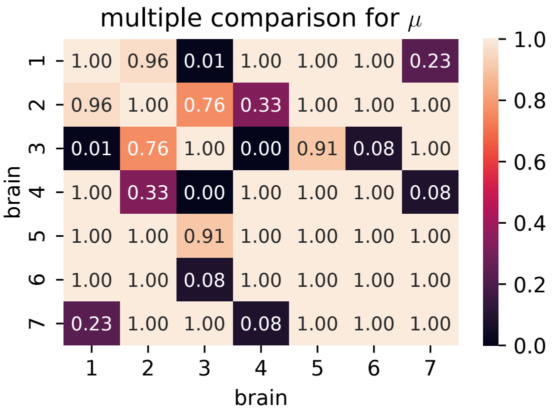


$\nu = 0.49$ and preconditioned data

Figure S8: Resulting $p$-values from pairwise post hoc Mann-Whitney-U tests comparing the shear modulus $\mu$ for different brains. Low $p$ values indicate that the samples from brain 3 show significantly different properties compared to those from the other tested brains.

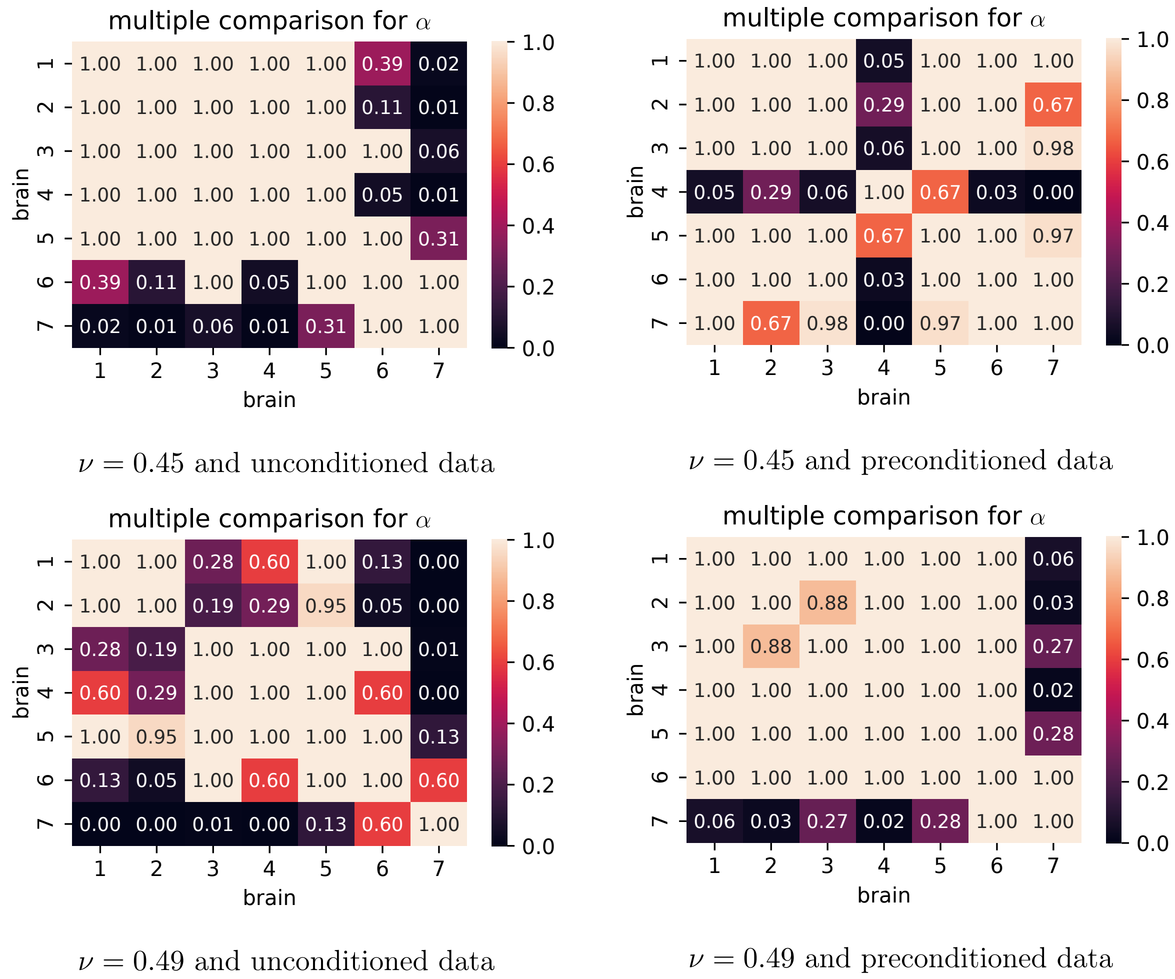


Figure S9: Resulting $p$-values from pairwise post hoc Mann-Whitney-U tests comparing the shear modulus $\mu$ for different brains. Low $p$ values show significantly different values for brain 7.

### 5 Influence of compressibility

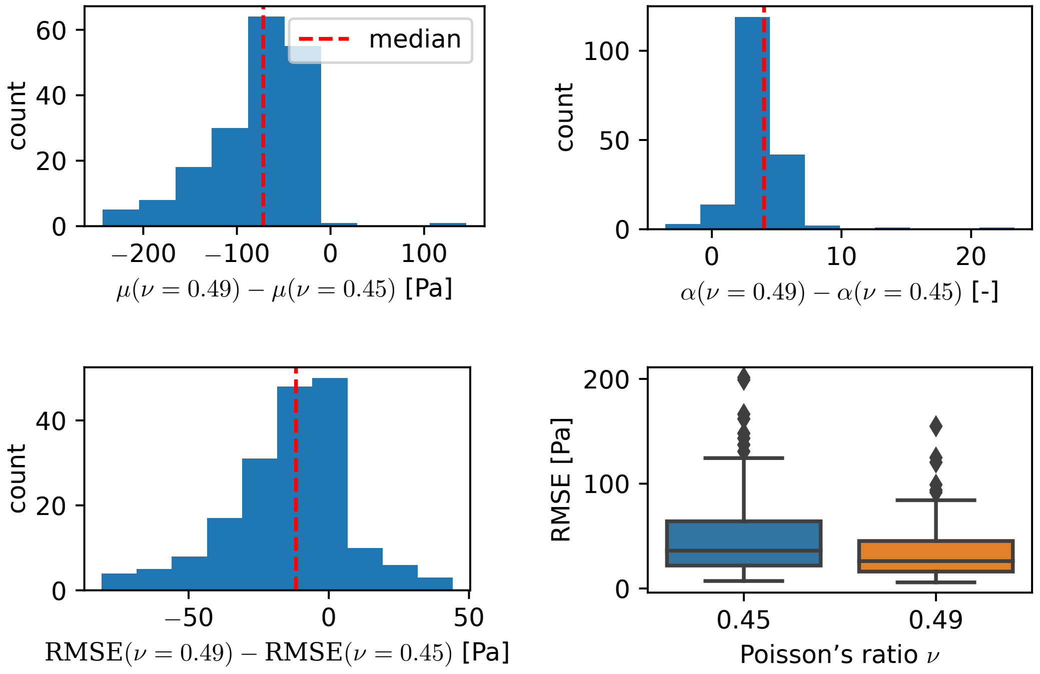


unconditioned data

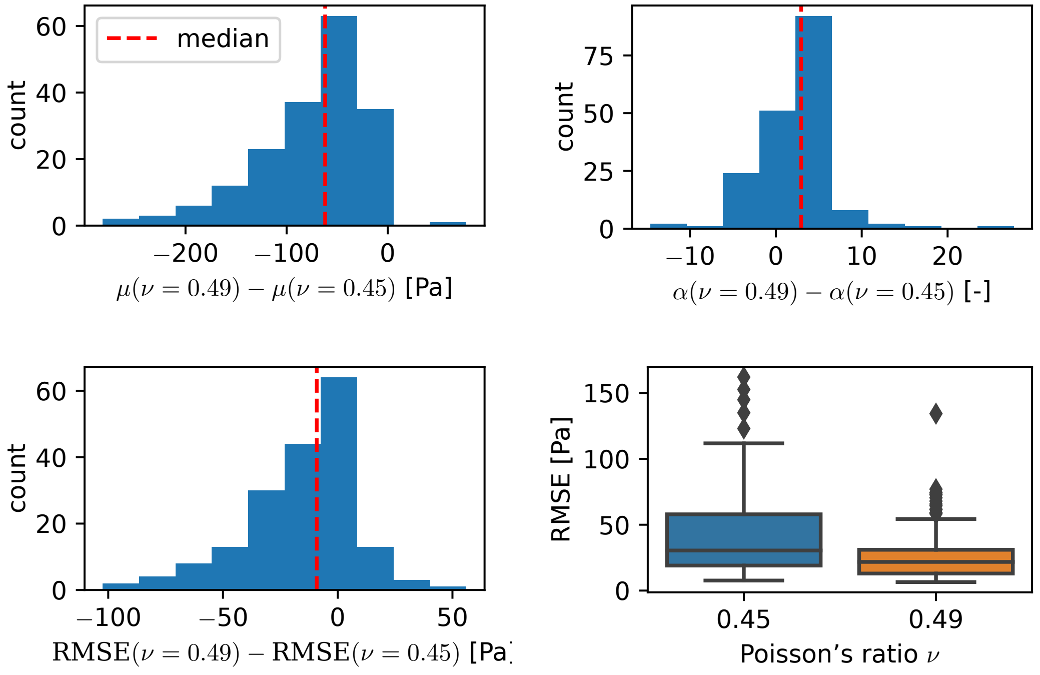


preconditioned data

Figure S10: Pairwise difference for the parameters $\mu$ and $\alpha$ and the RMSE between samples fitted with $\nu = 0.45$ and $\nu = 0.49$. The boxplots visualize the distribution of RMSE values to put the calculated differences in relation.

#### 6 Preconditioning behavior

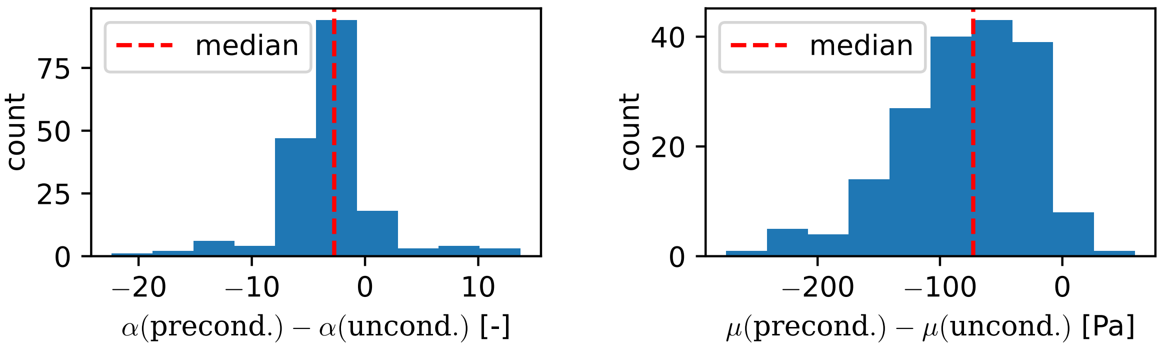


$\nu = 0.45$

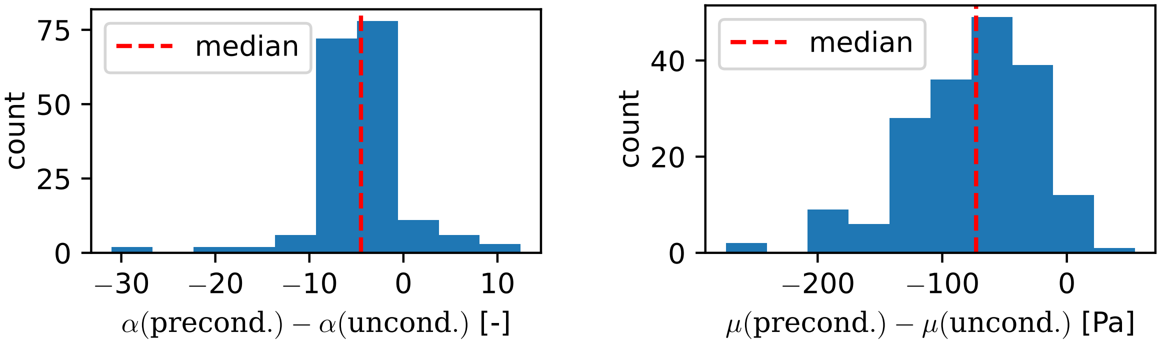


$\nu = 0.49$

Figure S11: Pairwise difference in the material parameters identified through fitting the un- and preconditioned data.

### 7 Averaging fitted parameters vs. fitting the averaged response

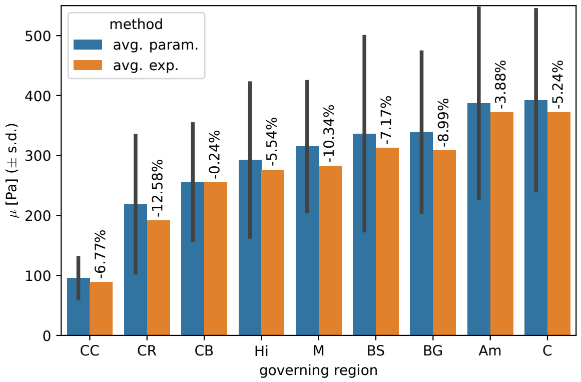


$\nu = 0.45$ and unconditioned data

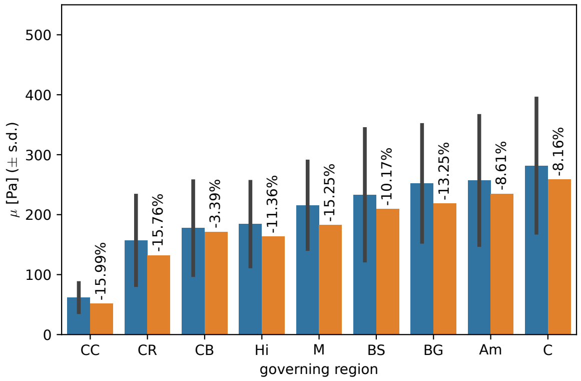


$\nu = 0.45$ and preconditioned data

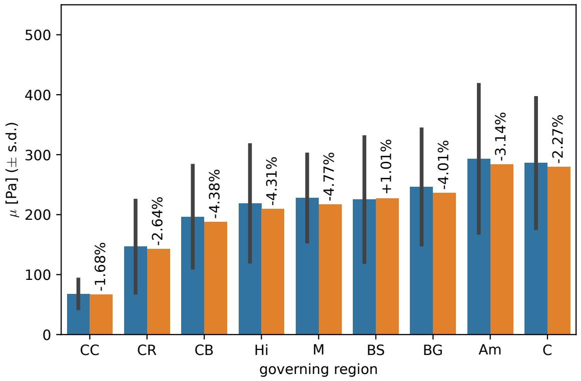


$\nu = 0.49$ and unconditioned data

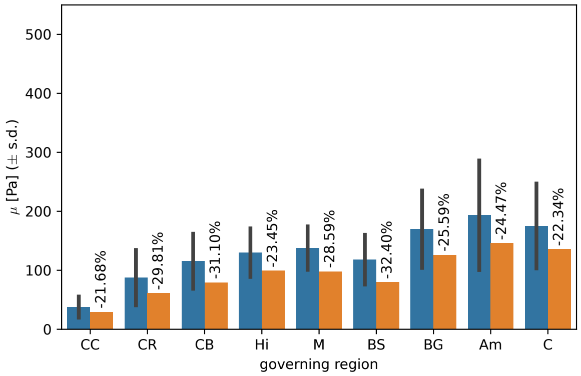


$\nu = 0.49$ and preconditioned data

Figure S12: Comparison between the averaged shear moduli $\mu$ obtained from fitting the experimental data of each specimen separately and those obtained from fitting the averaged experimental response.

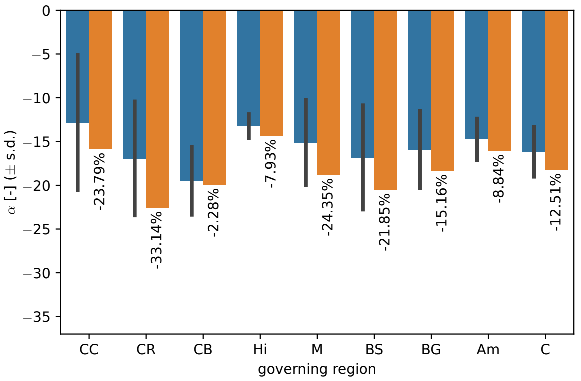


$\nu = 0.45$ and unconditioned data

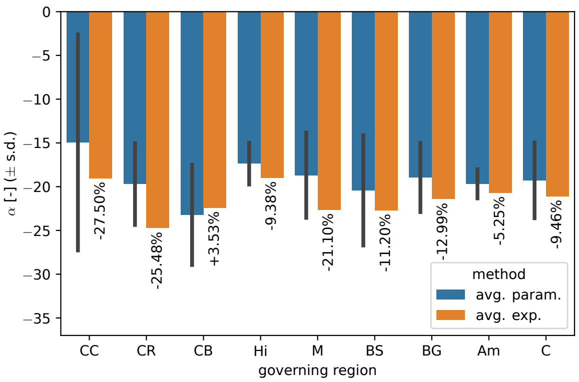


$\nu = 0.45$ and preconditioned data

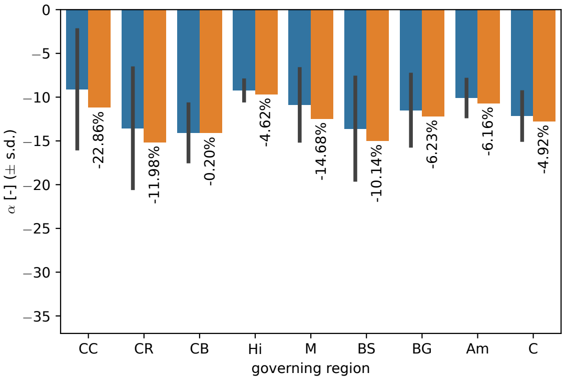


$\nu = 0.49$ and unconditioned data

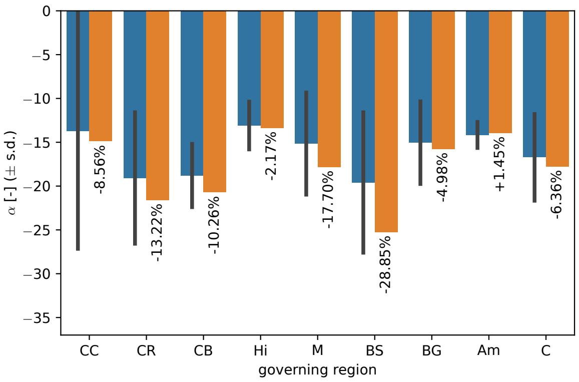


$\nu = 0.49$ and preconditioned data

Figure S13: Comparison between the nonlinearity parameter $\alpha$ obtained from fitting the experimental data of each specimen separately and those obtained from fitting the averaged experimental response.

### 8 Global optimality

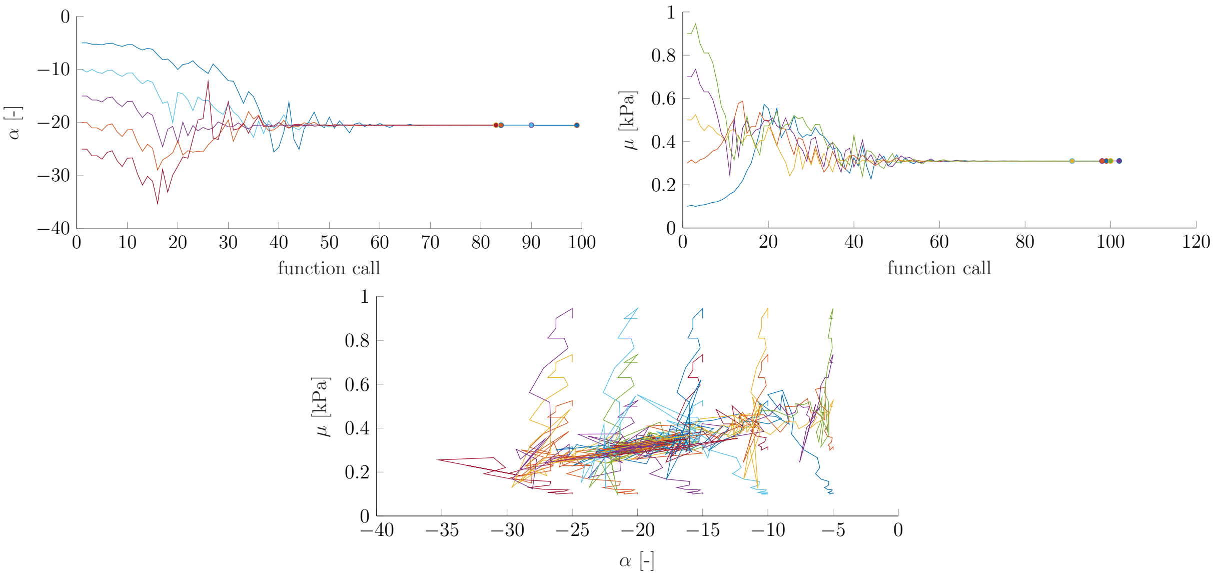


Figure S14: Global optimality study, where the optimization using the Nelder-Mead algorithm was started with different initial parameter values. The parameter values of the converged solution are close together, indicating no problems with local minima.
